# Single-Nucleus Multi-Omic Atlas Maps Regulatory Architecture and Non-Coding Variant Effects across Adult Human Tissues

**DOI:** 10.64898/2026.09.25.754561

**Authors:** Kaili Fan, Amrita Sule, Amy Guillaumet-Adkins, Ruochi Zhang, Tianxiong Yu, Zhijie Cao, Xuting Zhang, Neva Durand, MaryKate Tellier, Laura Domènech, François Aguet, Jason D. Buenrostro, Kristin G. Ardlie

**Author notes:** co-first authors (Kaili Fan, Amrita Sule). co-corresponding authors Correspondence: Kristin G. Ardlie, Jason D. Buenrostro, Kaili Fan.

## Abstract

Diverse human cell types establish specialized functions through lineage- and context-specific regulatory programs. Interpreting non-coding genetic risk requires integrated multi-omic reference maps that directly connect regulatory DNA to cellular expression across human tissues. Here we present a single-nucleus multi-omic atlas comprising 459,856 transcriptomic and chromatin accessibility profiles from 21 adult human tissues across four donors, including paired measurements from 160,688 nuclei. We resolve nine cell lineages, 61 broad cell types and 313 subclusters, and identify 1,085,062 candidate cis-regulatory elements (cCREs), including 161,270 novel elements absent from ENCODE. Regulatory activity was dominated by cell identity but refined by tissue context. Joint profiling enabled 871,177 cCRE-gene associations and revealed lineage-specific regulatory architectures. Cross-tissue analysis of accessibility further identified lineage-restricted and constitutively inaccessible chromatin domains, the latter showing preferential hypomethylation across human cancers. We leveraged this dataset to train sequence-to-function models to predict chromatin-accessibility effects for 548,656 fine-mapped variants, identifying 18,133 high-effect variants, including 1,120 broadly active variants. Models trained for eight endothelial subtypes further resolve predicted variant effects across vascular beds. Together, this study provides a comprehensive cellular and computational framework for interpreting regulatory sequence, context-dependent gene control, and complex trait genetics across the human body.

## Introduction

Diverse human cell types perform specialized functions despite sharing a largely common genome. Their identities are established by gene-regulatory programs in which transcription factors act on *cis*-regulatory elements to control gene expression and chromatin state. These programs reflect both developmental lineage and the tissue environments in which cells reside. Because most genetic risk for complex traits maps to non-coding DNA, understanding human physiology and disease requires determining which regulatory programs are conserved across the body, which are modified by local context, and how those programs connect regulatory sequence to cellular function^1–4^.

Single-cell transcriptomic atlases have transformed the classification of human cell types, while chromatin-accessibility maps have revealed the regulatory DNA that distinguishes them^3,5,6^. Joint measurement of RNA and chromatin accessibility connects transcriptional identity with regulatory state, providing complementary information for resolving closely related populations. Paired data also supports annotation when one measurement is sparse, and links accessible elements to gene expression within the same nuclei. Articulating this cellular diversity provides the context needed to interpret regulatory programs and predict where non-coding variants may act.

Biology is now moving from descriptive molecular catalogues toward predictive models that infer how DNA sequence and cellular context determine regulatory state and function. Recent sequence-to-function models predict gene expression, chromatin activity and effects of non-coding variation, while emerging AI virtual-cell frameworks seek to predict how cells respond to genetic and environmental perturbations^7–11^. Their biological scope, however, remains constrained by the measurements available for training and evaluation. Existing cross-tissue references frequently measure a single molecular layer, combine modalities generated in different samples or lack sufficient depth to resolve regulatory programs across primary adult cell types^3,5,12–16^. Predictive human biology therefore requires integrated, mechanistically grounded atlases that connect sequence, chromatin accessibility, transcription-factor activity and gene expression within defined cellular populations.

Here we generate an integrated RNA and chromatin accessibility atlas comprising 459,856 nuclei across 21 adult human tissues and four donors, including paired profiles from 160,688 nuclei^17^. The atlas resolves nine cell lineages, defined here as broad cellular compartments comprising related cell types. These lineages include 61 broad cell types and 313 Leiden-derived subclusters, which refine cell-type annotations and capture cellular heterogeneity. This hierarchy establishes a cellular framework for comparing shared regulatory programs and tissue-associated specialization. It defines 1,085,062 candidate *cis*-regulatory elements, including 161,270 absent from ENCODE^18^, links elements to genes and identifies lineage-specific and constitutively inaccessible domains. We then train sequence models across primary cell types and vascular endothelial subtypes to predict large accessibility effects of non-coding variants across traits and diseases. Together, these measurements and models connect cellular resolution to regulatory interpretation across adult human tissues.

## Results

### An integrated RNA and chromatin accessibility atlas resolves adult human cell types across tissues

We generated a single-cell ATAC-seq and RNA-seq (10x Genomics Multiome) atlas from 69 frozen samples previously collected and banked by the Genotype-Tissue Expression (GTEx) project in a collaboration with the ENCODE project^17^, spanning 21 adult human tissue sites and four donors, including two females and two males (**Fig. 1a**). The integrated design measured gene expression and chromatin accessibility from each captured nucleus, creating a cross-tissue reference in which regulatory and transcriptional states could be compared within a common experimental framework.

**Figure 1:**
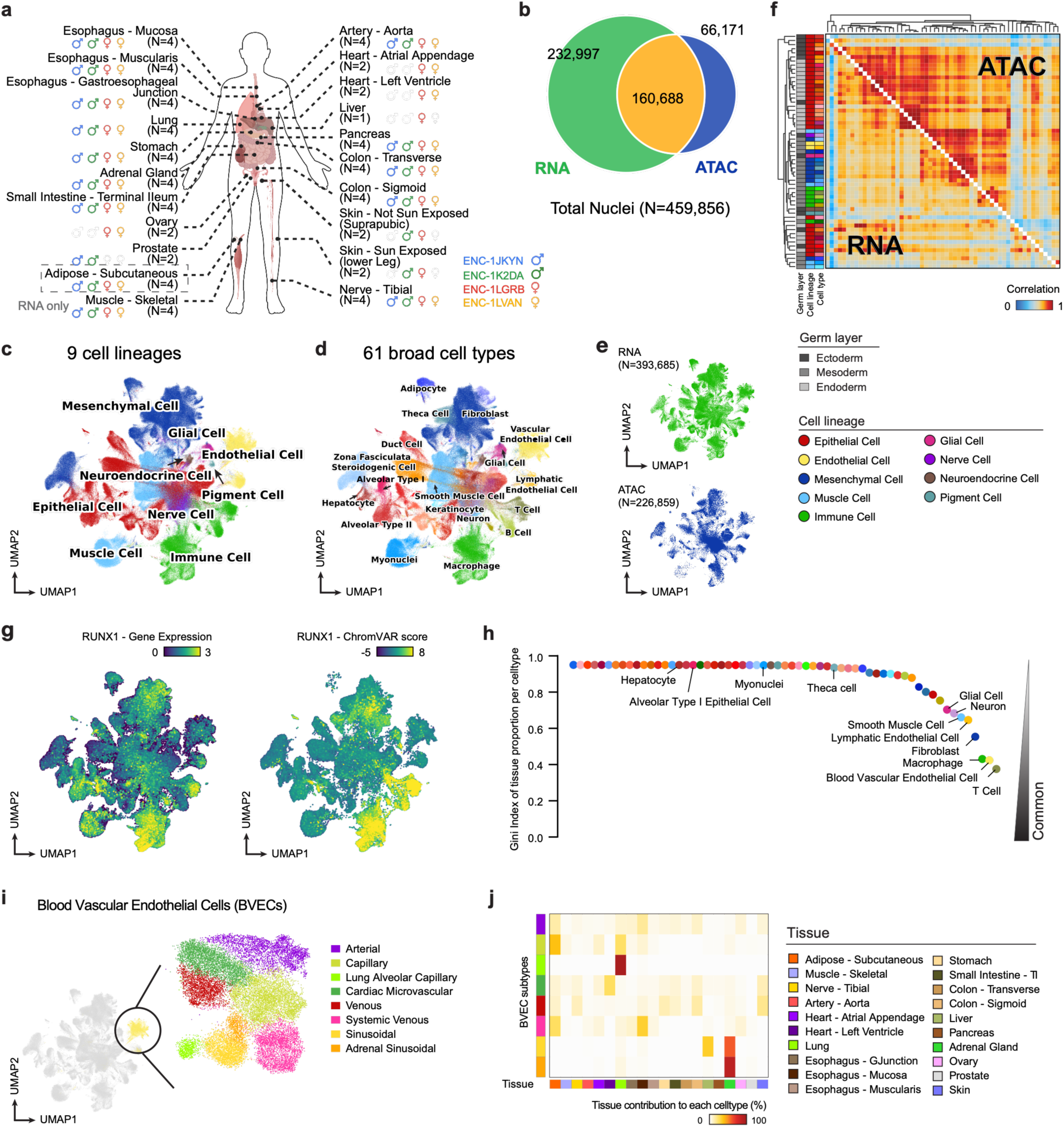
A cross-tissue 10x multi-omic atlas captures major cell types across adult human tissues. **a.** Schematic overview of the cross-tissue 10x Genomics Multiome dataset. Samples were collected from 21 adult human tissues contributed by four donors, including two males and two females, and spanning major anatomical systems. **b.** Venn diagram showing the numbers of nuclei profiled by RNA sequencing, ATAC sequencing or both modalities. **c.** UMAP embedding of all nuclei colored by nine major cell lineages. **d.** UMAP embedding of all nuclei colored by 61 broad cell types. **e.** UMAP embeddings of all nuclei with RNA profile (top) and ATAC profile (bottom), showing concordance. **f.** Cell-type similarity matrices based on chromatin accessibility (upper triangle) and RNA expression (lower triangle). Cell types were jointly ordered by hierarchical clustering of the integrated RNA–ATAC embedding, and the same order was applied to both modality-specific similarity matrices. Dendrograms and annotation bars indicate germ layer, cell lineage and broad cell type. **g.** UMAP embeddings colored by *RUNX1* expression (left) and *RUNX1* chromVAR motif-deviation score (right). **h.** Gini index of the proportion of each tissue that is composed of each broad cell type, highlighting common and tissue-restricted cell populations. **i.** UMAP embedding of endothelial nuclei highlighting blood vascular endothelial cells (BVECs) and colored by refined BVEC subtypes. **j.** Heatmap showing the percentage of nuclei in each endothelial subtype contributed by each tissue. The horizontal color bar below indicates tissue identity.

To generate this atlas, we optimized a nuclei-isolation protocol^19^ and developed a tissue- and sample-aware workflow for quality control, doublet detection, cross-sample integration and hierarchical annotation (**Methods**; **Supplementary Note 1; Supplementary Fig. 1**). Rather than applying uniform thresholds across organs with different cellular composition and data quality, the workflow combined orthogonal quality metrics and computational annotation with manual correction to reduce technical variation without removing rare or tissue-restricted cell populations.

RNA profiles outnumbered ATAC profiles, and incomplete overlap of the two modalities meant that excluding single-modality nuclei would reduce dataset size and rare-cell coverage. Many high-quality ATAC profiles also lacked sufficient RNA for direct paired annotation. We extended GLUE^20^ to transfer labels from confidently annotated nuclei with both modalities to RNA-poor nuclei using a joint representation of RNA and ATAC profiles. This strategy added annotation for an additional 66,171 RNA-poor nuclei, representing 14.4% of the atlas. Retaining these nuclei increased effective ATAC depth and representation of rare populations, extending the benefit of paired measurements to nuclei with incomplete RNA recovery.

The final atlas contained 459,856 high-quality nuclei. Of these, 393,685 (85.6%) had RNA profiles, 226,859 (49.3%) had ATAC profiles, and 160,688 (35.0%) had both modalities recovered (**Fig. 1b**). RNA profiles contained a median of 2,341 unique molecular identifiers (UMIs) and 1,374 detected genes, while ATAC profiles contained a median of 8,885 fragments and 10-fold transcription start site (TSS) enrichment across nuclei (**Extended Data Table 1**; **Supplementary Fig. 2a,b**).

Integrated RNA and chromatin accessibility profiles resolved nine major cell lineages and 61 broad cell types across the tissue–cell-type landscape. Within these annotations, Leiden clustering of the RNA profiles identified 313 subclusters, capturing finer transcriptional heterogeneity while preserving broad cell-type assignments. (**Fig. 1c–e**; **Extended Data Fig. 1a**; **Extended Data Table 2**). The atlas distinguished specialized tissue-populations within broad lineages, including adrenal zona fasciculata steroidogenic cells, ovarian theca cells and gastric foveolar epithelial cells, demonstrating the value of cross-tissue multi-omic profiling for refining cell-type annotation beyond broad lineage labels. Nuclei are grouped primarily by cell identity, while differences between RNA- and ATAC-based relationships capture complementary aspects of cellular organization (**Fig. 1f**). This annotation hierarchy established the cellular contexts for mapping regulatory variation across tissues.

Paired measurements also enabled comparison of TF transcript abundance with accessibility at DNA sequences containing the corresponding binding motifs, providing a comparative view of TF-associated regulatory activity across cell types. We quantified motif accessibility using chromVAR^21^ and related these scores to TF expression, following previously described approaches^22^. *RUNX1* expression was positively associated with motif accessibility across immune and stromal populations, whereas *FOXP1* showed weak or negative relationships in selected epithelial populations (**Fig. 1g; Extended Data Fig. 3; Methods**). Although transcript levels do not directly measure TF protein abundance, DNA binding or activity, these patterns suggest that TF transcript abundance alone does not consistently predict accessibility at associated motifs across cellular contexts.

The tissue distribution of cell types ranged from broadly shared fibroblast and vascular endothelial populations to organ-restricted parenchymal and epithelial populations^13,23^ (**Fig. 1h**; **Extended Data Fig. 1b–e**). We examined whether broadly distributed cell types could be resolved into populations with distinct tissue associations. Within blood vascular endothelial cells, integrated profiles distinguished eight subtypes, including lung alveolar capillary, cardiac microvascular and adrenal sinusoidal populations (**Fig. 1i,j**; **Extended Data Fig. 1f**). These subtypes provided an opportunity to test whether finer cellular resolution could reveal distinct predicted effects of disease-associated variants within a shared vascular lineage.

Compared with an independent single-cell ATAC atlas^5^ covering 20 overlapping adult tissues, the present atlas recovered concordant cell-type organization and additional sparsely sampled populations (**Extended Data Fig. 2a,b**). Across these 20 tissues, the present atlas contained 2.6 million total ATAC-seq fragments, compared with 1.6 million in the independent atlas, summed across all nuclei in each dataset. Paired RNA and ATAC measurements connected transcriptional annotations with regulatory profiles within the same nuclei, supporting a unified interpretation of cellular identity and regulatory state.

### Cell identity and tissue context organize the adult human regulatory landscape

We next used the cross-tissue accessibility profiles to define the regulatory elements underlying adult human cell identity. Aggregating ATAC signal within tissue–cell-type clusters produced 1,085,062 consensus candidate *cis*-regulatory elements (cCREs), represented as fixed 300-bp intervals covering 10.6% of the genome (**Supplementary Fig. 1**; **Supplementary Notes 1 and 2**). Most elements were TSS-distal, whereas promoter-associated elements were active across more cell clusters, consistent with broadly shared promoter activity and more context-dependent distal regulation^18,24,25^ (**Extended Data Fig. 4a–d**).

The atlas expanded existing catalogues of regulatory DNA. 323,450 elements were absent from an independent adult single-cell ATAC atlas^5^ (**Extended Data Fig. 2c**), and 161,270 were absent from ENCODE^18^ (**Fig. 2a**; **Extended Data Fig. 4e**). Across the atlas, 85.1% of atlas cCREs overlapped ENCODE, with elements absent from ENCODE concentrated in rare or specialized cell types (**Extended Data Fig. 4c**), highlighting the contribution of cellular resolution to regulatory discovery. Of the 3,889 experimentally validated VISTA enhancers^26^, 2,353 (59.7%) overlapped atlas cCREs. Conversely, 5,657 atlas cCREs overlapped VISTA enhancers, including 298 cCREs that were not represented in ENCODE, providing independent support for their regulatory activity.

**Figure 2:**
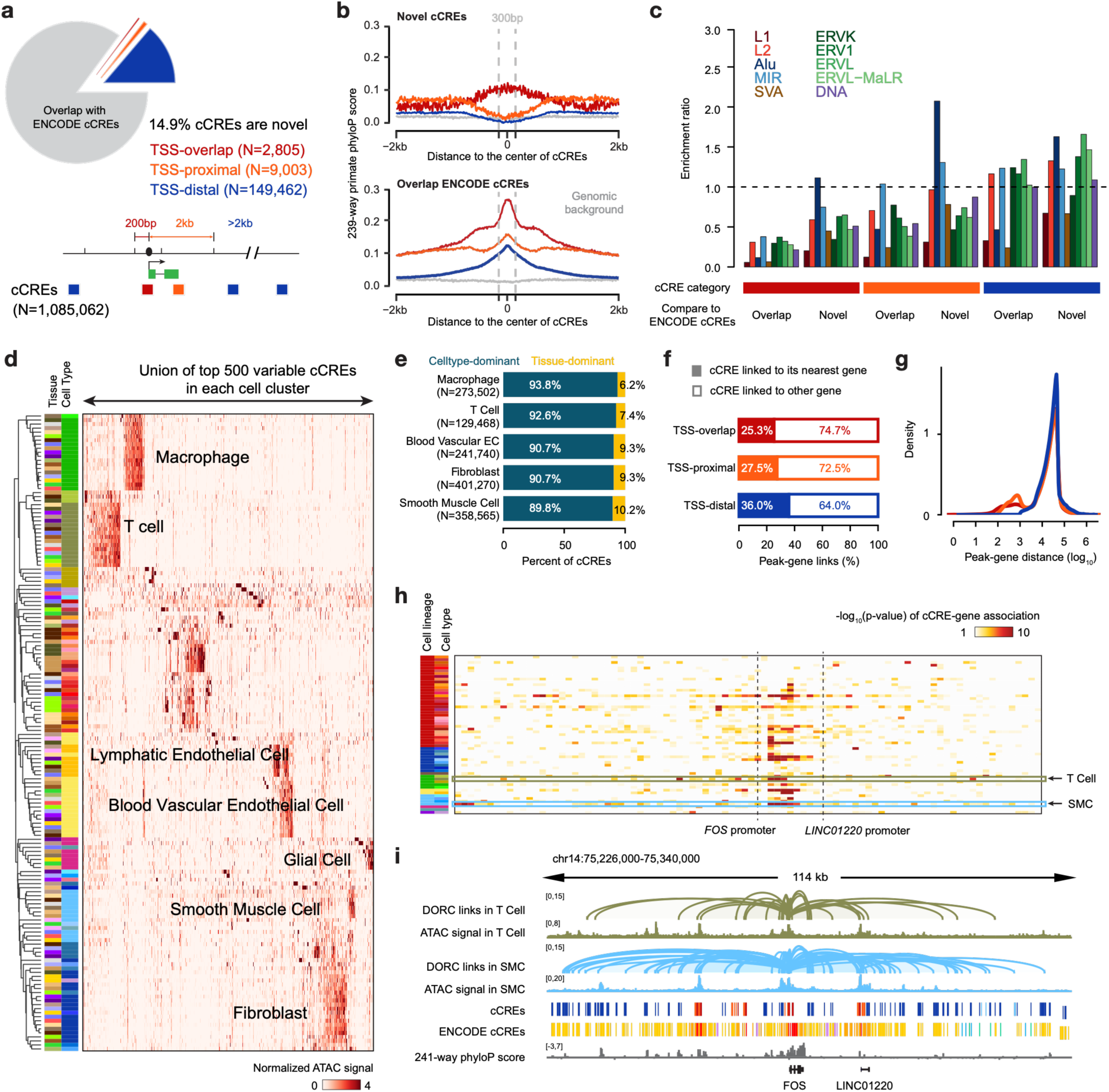
A cross-tissue atlas of candidate *cis*-regulatory elements in adult human cell types. **a.** Classification of cCREs according to overlap with the ENCODE cCRE catalogue. The pie chart shows ENCODE-overlapping cCREs and novel cCREs; the inset lists the numbers of novel cCREs in each genomic-context group. **b**. Aggregation plot of 239-way primate phyloP scores centred on cCREs, stratified by genomic-context group and by overlap with the ENCODE cCRE catalogue. **c.** Enrichment ratio of cCREs with annotated transposable-element families, stratified by genomic-context group and cCRE novelty. The dashed line indicates the expected value of 1. **d**. Heatmap showing normalized ATAC-seq signal for the union of the 500 most variable cCREs identified in each cell cluster. Rows represent cell clusters ordered by hierarchical clustering, and columns represent cCREs. Cell-type annotations are indicated alongside the heatmap. **e.** Bar plot showing the percentage of cCREs classified as cell-type-dominant or tissue-dominant across five common cell types. **f**. Percentage of peak-gene links connecting cCREs to their nearest gene or to another gene, stratified by genomic-context group. **g**. Density distribution of peak-gene distances, shown on a log_10_ scale. **h**. Heatmap of cCRE-gene association *p*-values for cCREs linked to *FOS* across cell types. The annotation bars alongside the heatmap indicate cell lineage and cell type. cCREs are ordered by genomic position to match the browser view in i. **i**. Genome-browser view of the *FOS* and *LINC01220* loci in T cells and smooth muscle cells (SMC), showing DORC links, ATAC-seq signal, and 241-way phyloP scores. Track colors identify the cell type, whereas signal magnitude is represented by track height. In the cCRE and ENCODE cCRE tracks, colors indicate cCRE categories as in Fig. 2a and **Extended Data Fig. 4e**.

Distal cCREs absent from ENCODE showed weaker evolutionary constraint and greater cell-type specificity than previously annotated elements. They were enriched for sequence classes with limited alignment across mammals (**Extended Data Fig. 5**), while negative phyloP scores supported accelerated evolution of a subset of elements across primate species (**Fig. 2b**; **Extended Data Fig. 4f–h**). Rapidly evolving cCREs were enriched in keratinocytes, melanocytes, colonocytes and hepatocytes, implicating barrier, detoxification and metabolic cell types in regulatory innovation (**Extended Data Fig. 4k**). These observations distinguish newly catalogued regulatory DNA from previously annotated elements and motivated analysis of the contribution of transposable elements.

Transposable elements contributed prominently to this recently evolved regulatory sequence. TEs overlapped 54.6% of novel cCREs, compared with 31.4% of ENCODE-overlapping cCREs, with enrichment for Alu and LINE2 families (**Fig. 2c**; **Extended Data Fig. 4j**; **Supplementary Fig. 3**). Although these elements were weakly conserved overall (**Extended Data Fig. 4i,5c**), localized constraint and recurrent positional hotspots were consistent with selective retention of functional motifs within the co-opted TE sequence.

To distinguish regulatory programs shared across organs from tissue-associated adaptations, we compared accessibility in five broadly distributed cell types. Among cCREs classified according to whether accessibility was driven primarily by cell type or tissue, 89.8–93.8% of cCREs were classified as cell-type-dominant and 6.2– 10.2% as tissue-dominant (**Fig. 2d,e**; **Extended Data Fig. 6a,b**; **Methods**). Shared lineages therefore retained a conserved regulatory structure across organs, with a smaller set of elements capturing tissue-specific variation.

Newly catalogued elements were frequently associated with gene expression. Overall, 36.9% of cCREs absent from ENCODE were linked to at least one gene; among distal elements, the proportion was similar for novel and previously annotated cCREs (42.4% versus 43.8%). A novel distal element associated with *PROX1* expression in pancreatic vascular endothelial cells illustrates how the expanded catalogue nominates cell-type-restricted regulatory relationships^27,28^ (**Extended Data Fig. 4l**).

### Paired RNA and chromatin resolve cell-type-specific regulatory circuits

To connect accessible elements with transcriptional output, we developed fastDORC to identify elements whose accessibility covaries with gene expression across nuclei. This scalable implementation of the Domains of Regulatory Chromatin (DORC) framework^29^ models technical covariates and supports analysis of rare populations (**Supplementary Notes 3**). On identical inputs, fastDORC was approximately 116-fold faster than the original DORC implementation (12 min versus 23.16 h) while recovering 24,369 shared cCRE–gene associations and comparable association rankings (Spearman’s rho = 0.762; **Supplementary Note 3; Supplementary Fig. 4a,b**). The remaining differences likely reflect fastDORC’s optimized statistical framework, including its treatment of technical covariates and sparse or rare cell populations, rather than differences in the underlying input data. Applied independently across cell types, fastDORC identified 871,177 cCRE–gene associations involving 19,459 genes, with a median of 55,315 links per cell type (**Extended Data Fig. 6d; Supplementary Note 4**).

DORC activity, defined as the aggregate accessibility of cCREs linked to a gene, recapitulated gene-expression patterns^29^. For example, DORC activity and RNA expression showed concordant patterns for *VWF* (Von Willebrand Factor) across vascular populations^30^ (**Extended Data Fig. 6c**). Overall, 47.3% of cCREs were linked to at least one gene, with a median peak–gene distance of approximately 10 kb, while 36.0% of distal cCRE– gene links involved a non-nearest gene (**Fig. 2f,g**). Thus, genomic proximity alone would miss a substantial fraction of the regulatory associations identified by paired measurements.

We next identified genes associated with unusually large numbers of regulatory elements, defining DORC genes as those with more than twice the mean number of cCRE–gene links per gene within a cell type (**Methods**). The atlas contained 11,492 DORC genes, with a median breadth of four cell types (**Extended Data Fig. 6d,e**). Broadly shared DORCs were enriched for vascular, migratory, haemostatic and cytoskeletal functions, whereas restricted DORCs captured transcriptional, metabolic, stress-response, cell-cycle and immune programs (**Extended Data Fig. 6f,g**).

Even when a target gene was shared, its associated regulatory elements differed across cellular contexts. FOS-linked cCREs varied in accessibility and association strength across cell types, with T cells and smooth muscle cells showing distinct associated element repertoires around the same locus (**Fig. 2h,i**). Thus, a shared gene can be associated with different combinations of regulatory elements in different cell types, consistent with the regulatory billboard model of context-dependent enhancer function^31,32^.

Together, the tissue-aware processing workflow, GLUE-based recovery of RNA-poor nuclei and fastDORC provide an atlas-scale toolkit for integrated RNA and chromatin accessibility data. The resulting maps reveal that cell identity depends not only on expressed genes but also on context-specific cCRE combinations. Shared genes can therefore be regulated through distinct circuits that support conserved identity and tissue specialization.

### Cross-tissue accessibility identifies lineage-specific and constitutively inaccessible domains

The breadth of the atlas allowed us to examine regulatory organization above the scale of individual cCREs. We asked whether genomic regions inaccessible across diverse cell types define a shared heterochromatin state, distinct from domains accessible in selected lineages. Cross-tissue accessibility provided the map for this comparison, while independent chromatin measurements allowed us to test its correspondence with biochemically defined heterochromatin^33,34^.

We partitioned the genome into 26,502 non-overlapping 100-kb windows, excluding regions with low mappability (**Methods**), and quantified ATAC signal and cCRE activity across tissue–cell-type clusters. This scale captures broad domains while retaining resolution for comparisons across cellular contexts^34^. Unsupervised clustering separated ATAC-enriched regions (AERs), cell-type-specific ATAC-depleted regions (cts-ADRs) and ubiquitously ATAC-depleted regions (ubi-ADRs), here termed constitutively inaccessible domains (**Fig. 3a,b**; **Extended Data Fig. 7a**).

**Figure 3:**
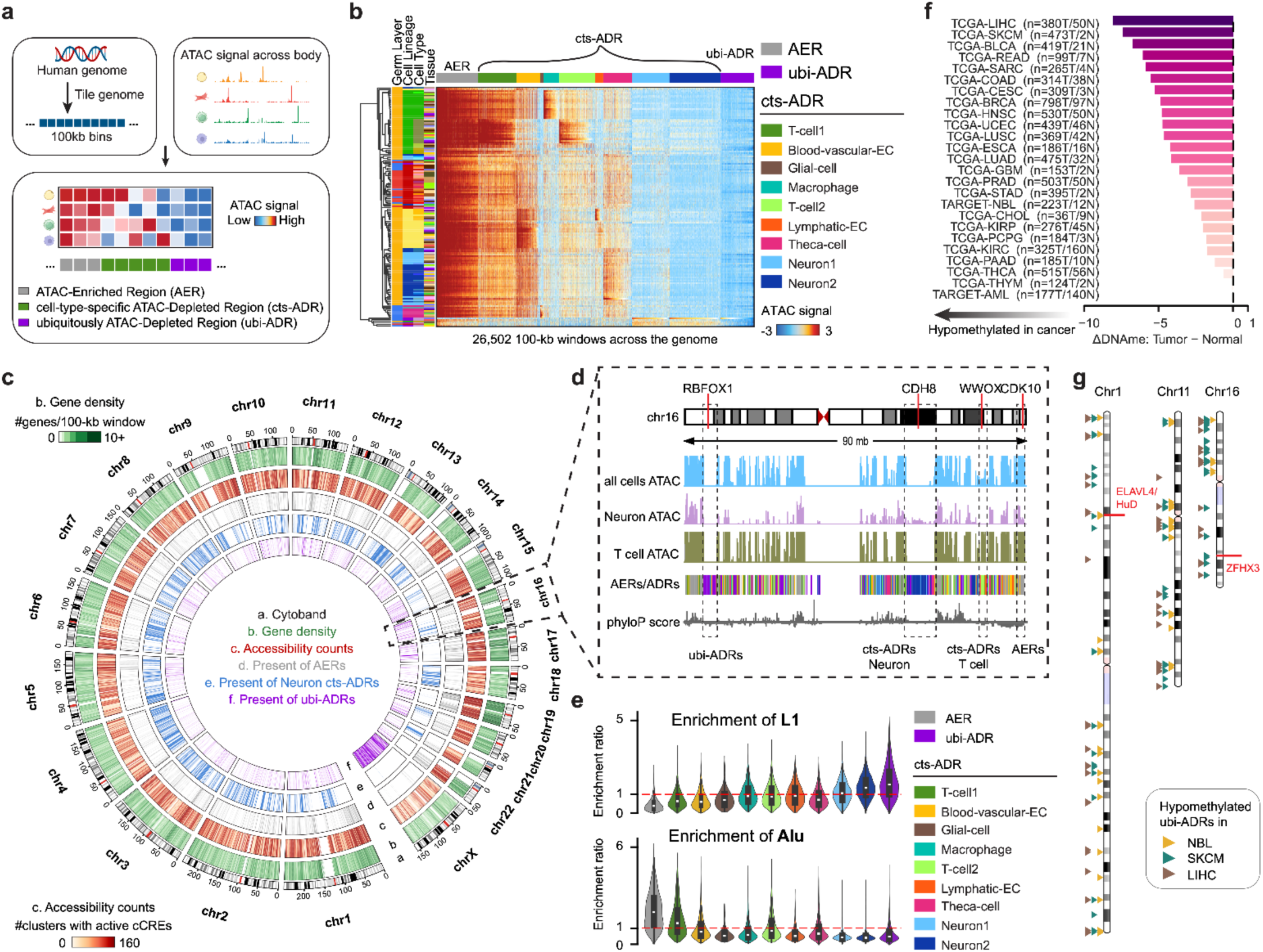
Genomic landscape of broad chromatin domains across human cell types. **a.** Overview of the accessibility-domain analysis. The genome was divided into 100-kb windows and classified by shared accessibility patterns across cell types. See **Methods** for details. **b.** Accessibility across genomic windows and cell types. Rows represent cell clusters and columns represent genomic bins ordered by hierarchical clustering. Annotation bars indicate germ layer, cell lineage, cell type and tissue of origin. The bar above the heatmap denotes AERs, cts-ADRs and ubi-ADRs; colors indicate the corresponding domain subclasses. **c**. Genomic distribution of domain classes. **d.** Representative view of chr16 highlighting broadly active, lineage-specific and constitutively inaccessible domains. **e.** L1 and Alu enrichment across domain classes. **f**. Cancer-associated DNA methylation changes across domains. **g**. Constitutively inaccessible domains hypomethylated in neuroblastoma (NBL), melanoma (SKCM) and liver cancer (LIHC).

AERs were accessible across most tissues and cell types, whereas cts-ADRs were inaccessible in most populations but accessible in selected lineages. Distinct cts-ADR subclasses showed accessibility in T cells, vascular and lymphatic endothelial cells, macrophages, glia and theca cells and were enriched for functions appropriate to those populations (**Fig. 3b**; **Extended Data Fig. 7b**). These domains capture regulatory organization across extended genomic intervals beyond individual accessible elements.

To test whether constitutively inaccessible domains correspond to heterochromatin, we compared them with independent chromatin measurements. These domains comprised approximately 10.2% of the genome, containing few active cCREs (**Extended Data Fig. 7a**), and were enriched for the repressive histone modifications H3K9me3 and H3K27me3^35^ (**Extended Data Fig. 7c**). Critically, 79.3% overlapped sonication-resistant heterochromatins (srHCs) previously mapped in fibroblasts, providing orthogonal biochemical validation of their compacted heterochromatic state^34^. Together, these observations support the conclusion that cross-tissue inaccessibility identifies a constitutive heterochromatin compartment, connecting a shared pattern of inaccessibility across human cell types to an independently measured physical property of chromatin.

Representative loci illustrated the distinction between broad and lineage-selective accessibility. *CDH8*^36^ overlapped a neuron-active cts-ADR, whereas the broadly expressed *CDK10* locus fell within an AER (**Fig. 3d**). The X chromosome likewise contained extensive ubi-ADRs interrupted by accessible domains, including an AER at *DDX3X*, an X-inactivation escape gene^37^ (**Extended Data Fig. 7g**). These examples show how domain-level analysis separates constitutive repression from regulatory neighborhoods selectively available in particular cellular contexts.

The three domain classes occupied distinct sequence environments. Repressive classes occurred preferentially in gene-poor, AT-rich and weakly conserved regions, with ubi-ADRs showing the lowest C+G content and CpG density (**Fig. 3c**; **Extended Data Fig. 7d,e**). ubi-ADRs were also enriched for AT-rich LINE1 elements and depleted of GC-rich Alu and LINE2 elements (**Fig. 3e**; **Extended Data Fig. 7f**), contrasting with the Alu- and LINE2-enriched sequences found among active novel cCREs. These intrinsic genomic features may contribute to the distinct epigenetic behavior of constitutively inaccessible domains in disease.

We next asked whether domains consistently inaccessible in normal tissues are preferential sites of cancer-associated DNA methylation loss. Across 9,051 tumour and tissue-normal methylation profiles from 25 TCGA and TARGET projects, ubi-ADRs showed the greater tumour-associated hypomethylation than other domains in 23 of 25 projects (one-sided Mann–Whitney U, FDR < 0.05), with losses reaching Δβ = −0.08 in the most affected cancers. By contrast, AERs and cts-ADRs showed smaller and more heterogeneous changes^38–41^ (**Fig. 3f**; **Extended Data Fig. 8a**). Different tumour types showed methylation loss in partially distinct subsets of domains, including regions near *ELAVL4* in neuroblastoma and *ZFHX3* in melanoma^42–44^ (**Fig. 3g**; **Extended Data Fig. 8b,c**). Thus, normal cross-tissue accessibility identifies genomic compartments with distinct patterns of cancer-associated methylation loss.

Together, these results define an accessibility-based hierarchy that ranges from broadly active regions to lineage-selective and constitutively inaccessible domains. Their sequence composition, overlap with sonication-resistant heterochromatin, and preferential tumour-associated hypomethylation support a stable heterochromatin compartment that is remodeled in disease, although the functional consequences of that remodeling remain to be established.

### Sequence models predict cell-type-specific effects of genetic variation

Having mapped regulatory organization across tissues, we next asked how disease-associated sequence variants might alter accessibility in these cellular contexts. Advances in sequence-to-function modeling now support prediction of molecular activity and non-coding variant effects from DNA sequence^7–10^. We selected seq2PRINT^7^ because it predicts chromatin-accessibility and transcription-factor-footprint profiles at base resolution, while its scalable architecture supports training across many primary cell types and evaluation of hundreds of thousands of variants.

Before applying seq2PRINT across traits and diseases, we validated its predictions against independently measured chromatin-accessibility QTL effects in monocytes, B cells, CD4 and CD8 T cells. We trained seq2PRINT using bulk and single-nucleus ATAC-seq data from Corces et al.^45^ and ENCODE portal^35^, and evaluated its predictions against caQTL effects from the CIMA atlas of 428 individuals^15^. Predicted and measured effects were strongly correlated on all datasets, reaching Pearson r ≈ 0.73 in monocytes and CD4 T cells, r=0.79 in CD8 T cells and r=0.8 in B cells at highest training depths examined (**Extended Data Fig. 9a-b**). Because many cell types in our atlas contained fewer than 10 million unique fragments, we next asked how training depth influences variant-effect prediction. Downsampling showed that correlations generally improve with depth, with 10 million fragments yielding performance close to that obtained at the highest depths (r ≈ 0.73 and 0.73 for CD4 T cells and monocytes, respectively). Models trained with only 2 million fragments retained substantial predictive power (r ≈ 0.58 and 0.59, respectively) (**Extended Data Fig. 9c-d**). We therefore selected a threshold of 10 million fragments to balance predictive performance with cellular coverage, training models for 35 primary cell types. The performance retained at lower depths suggests that modeling could extend to 49 cell types represented by at least 2 million fragments. Together, these results validate seq2PRINT variant-effect predictions in primary immune cells and support extending this approach to more finely resolved cellular populations.

We evaluated the resulting models on held-out chromatin-accessibility and TF-footprint profiles, confirming their ability to recover cell-type-specific regulatory activity (**Supplementary Fig. 5**). We next extended variant prediction across traits and diseases, assembling 588,618 variants from fine-mapped credible sets (groups of candidate causal variants) across 48,828 Open Targets studies^46^. These credible sets were generated using source- and data-dependent fine-mapping approaches, including SuSiE^47^, SuSiE-inf^48^ and PICS^49^. The posterior inclusion probability (PIP) represents the model-based probability that a variant is causal for an association, conditional on the data, linkage disequilibrium structure and assumptions of the relevant fine-mapping method. Because PIP calibration may vary across studies (**Supplementary Fig. 6a**), we therefore evaluated PIP thresholds from 0.1 to 1.0 and selected one threshold per study using the median enrichment of predicted regulatory effects across cell types (**Supplementary Note 5**; **Supplementary Fig. 6b-c**). After quality control, the framework evaluated 548,656 SNPs across 6,631 studies, asking whether trait-associated variants were enriched for predicted regulatory effects in specific cell types relative to variants from other studies matched for cell type and PIP threshold (**Fig. 4a**; **Supplementary Note 5**).

**Figure 4:**
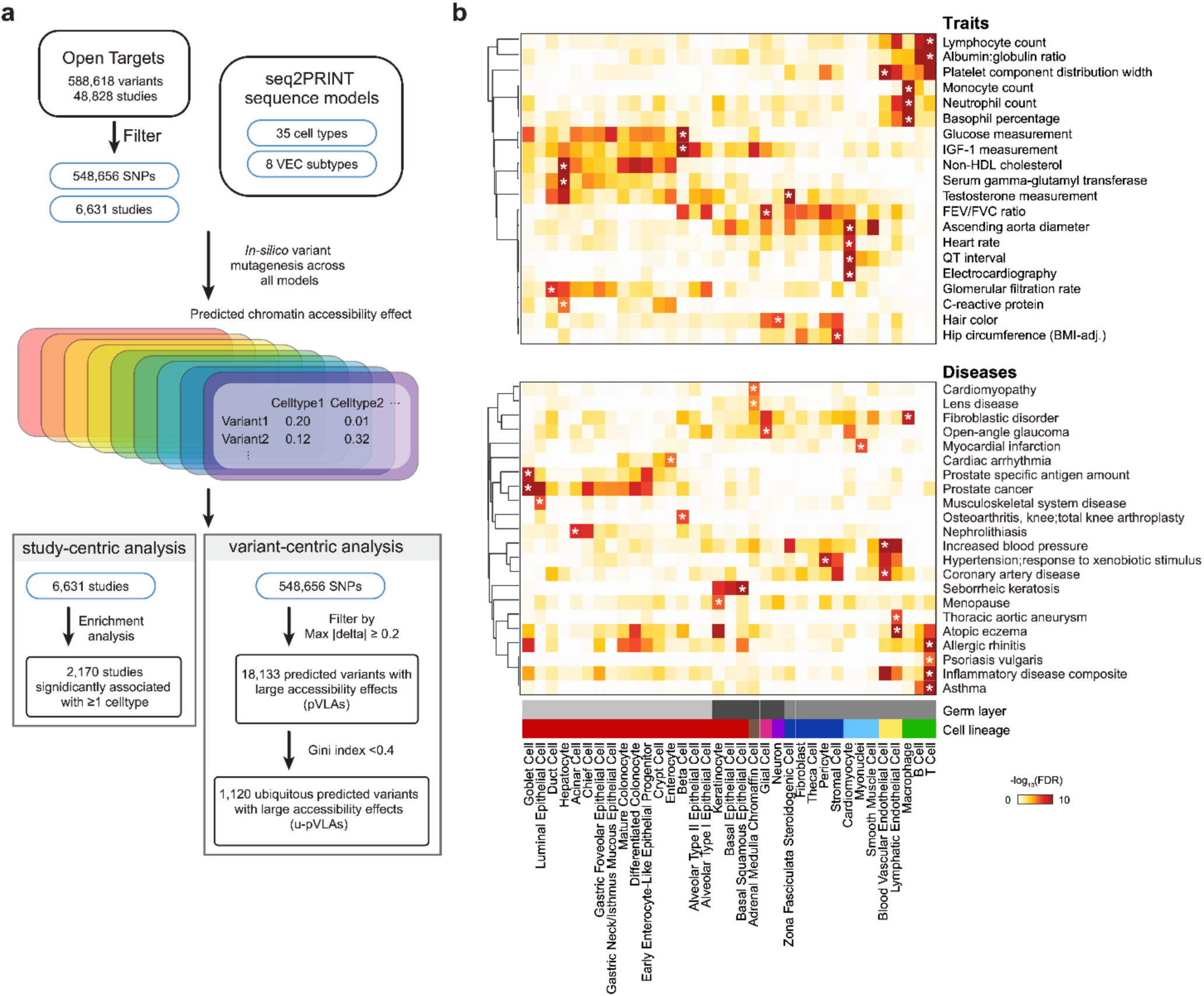
Cell-type-resolved regulatory architecture of complex traits using seq2PRINT. **a.** Overview of the seq2PRINT analysis pipeline. seq2PRINT models trained for 35 primary cell types and 8 blood vascular endothelial cell subtypes were applied in silico to variants in Open Targets fine-mapped credible sets. Of 588,618 variants across 48,828 GWAS studies, 548,656 variants from 6,631 studies were retained after quality control, credible-set filtering and study-specific PIP-threshold optimization. Variant-level predicted regulatory effects were aggregated by study and cell type, identifying 2,170 studies with significant associations in at least one cell type. For variant-centric analysis, variants with a maximum absolute predicted effect of at least 0.2 were classified as predicted variants with large accessibility effects (pVLAs). Variants with a Gini coefficient below 0.4 across the 35 primary cell types were classified as ubiquitous pVLAs (u-pVLAs). **b**. Heatmap showing associations between selected traits and diseases and the 35 primary cell types. Colors indicate −log_10_(FDR), and annotation bars indicate germ layer, cell lineage. Asterisk represents the cell type–trait/disease pair with the best score for each trait/disease.

Study-level predicted effects recovered biologically coherent relationships between traits and cellular context. Among 2,170 studies with a significant association in at least one model, blood-cell traits mapped to corresponding immune and myeloid populations, whereas glucose, IGF-1 and lipid traits preferentially mapped to hepatocytes and adipocytes. Cardiovascular phenotypes-including myocardial infarction, cardiomyopathy and coronary artery disease-were enriched in cardiomyocyte and vascular models, while asthma and allergic disease implicated both epithelial and immune cell types (**Fig. 4b**; **Extended Data Fig. 9e**). Predicted-effect profiles further separated related populations more clearly than raw accessibility alone, indicating that the models capture regulatory information not apparent from accessibility magnitude alone (**Extended Data Fig. 9f**). Associations extending beyond canonical disease tissues may reflect shared regulatory programs, pleiotropic variants, activity in multiple cell types or limitations of the current models^46^. We therefore interpret cell-type associations and variant-effect scores as predictions that prioritize cellular contexts and mechanisms for experimental investigation, rather than as evidence of disease causality.

Together, these analyses connect integrated primary-cell training data with scalable variant prediction and study-specific selection of fine-mapped variants. The resulting map recovers expected trait–cell-type relationships and nominates additional cellular contexts across thousands of phenotypes, providing a framework for investigating regulatory mechanisms beyond a single organ or lineage.

### Predicted variant effects range from cell-type-restricted to broadly active

We next examined individual variants with large predicted allelic effects, measured as the difference in predicted accessibility between alternate and reference alleles. Across 548,656 modeled SNPs, 18,133 (3.3%) exceeded an absolute effect-score threshold of 0.2 in at least one of 35 cell-type models and were designated predicted variants with large accessibility effects (pVLAs; **Fig. 5a**; **Methods**). The score is expressed as a unitless difference in model-predicted accessibility on the model’s normalized output scale. Larger absolute values indicate larger predicted allelic effects, whereas the sign indicates direction: positive values predict increased accessibility for the alternate allele, with negative values predicting decreased accessibility. The threshold of 0.2 was used as an operational definition of a large predicted allelic effect and was not calibrated to a specific fold change or experimentally measured difference in accessibility. Of these variants, 52.7% overlapped atlas cCREs used for model training, while the remainder overlapped external cCRE annotations or fell outside current catalogues (**Fig. 5b**).

**Figure 5:**
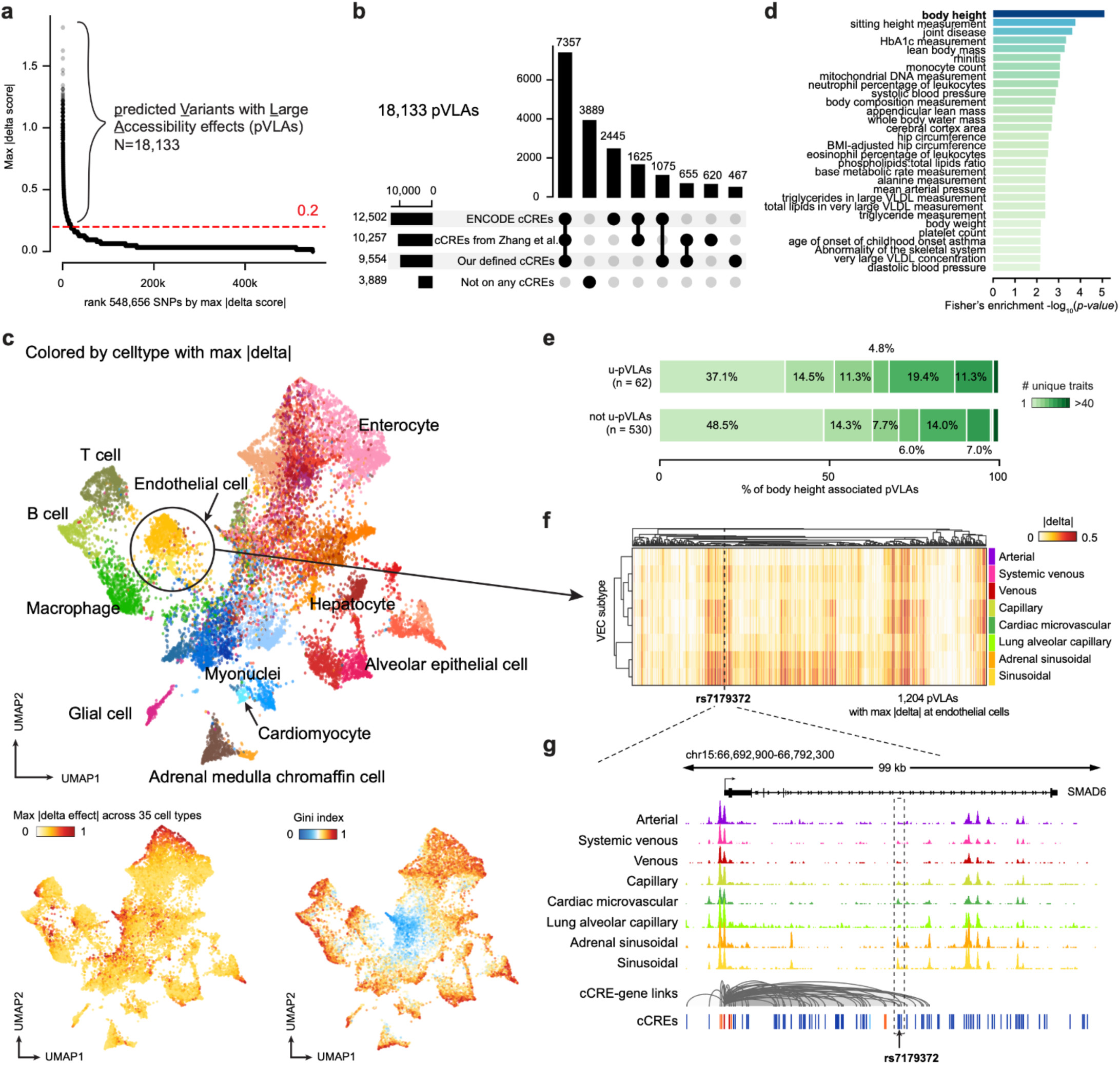
Cell-type-specific regulatory effects of complex-trait-associated variants. **a.** Distribution of the maximum absolute predicted regulatory effect across variants. Variants are ranked according to their maximum predicted effect across the 35 primary cell types; the dashed red line indicates the selection threshold of 0.2. The number of pVLAs exceeding this threshold is indicated. **b**. Overlap of pVLAs with cCREs defined in this study, cCREs reported by Zhang *et al.* and the ENCODE cCRE catalogue. **c**. UMAP embeddings of variant regulatory-effect profiles across the 35 primary cell types. Bottom Left, variants colored according to the cell type in which their maximum absolute predicted effect occurs. Upper, the same embedding annotated by representative cell types and lineages. Bottom Right, variants colored by the Gini coefficient of their absolute predicted-effect profiles, with blue indicates gini score<0.4. **d**. Traits enriched for ubiquitous pVLAs (u-pVLAs). The x axis shows the Fisher enrichment −log_10_(*p-value*). **e**. Pleiotropy of u-pVLAs and non-u-pVLAs across 6,631 Open Targets GWAS studies. Bars show the percentage of variants associated with one or more traits. **f.** Heatmap of predicted regulatory effects of pVLAs associated with blood vascular endothelial cell subtypes. Rows represent endothelial subtypes and columns represent pVLAs with max effect size identified in endothelial cells. **g.** Genome-browser view of the *SMAD6* locus showing ATAC-seq signal across blood vascular endothelial subtypes, cCRE-gene links, cCREs and the position of rs7179372.

The predicted effects formed a continuum from restricted to broadly distributed activity. Variant-effect profiles clustered according to the cell type in which the largest effect occurred, with related immune, vascular, epithelial and parenchymal populations occupying neighboring regions (**Fig. 5c**). The UMAP-defined clusters showed enrichment for lineage-associated TF binding in independent ChIP-seq data^50^, providing orthogonal support that the predicted effects reflect underlying regulatory programs (**Extended Data Fig. 10a,b**).

A subset of 1,120 variants (6.2%) had predicted effects that were broadly distributed across cell types and were classified as ubiquitous pVLAs (u-pVLAs), defined as those with a Gini coefficient below 0.4, indicating less individual cell type effects (**Fig. 5c**). Within this u-pVLA subset, 1,076 variants had assigned domain annotations, with 406 (37.7%) overlapping AERs, compared with 13.0% genome-wide, corresponding to an approximately 2.9-fold enrichment (Fisher’s exact test, *p-value*=1×10^-100^). These variants were enriched among highly polygenic traits, with body height showing the strongest enrichment (*p-value*=8×10^-6^, **Fig. 5d**). Among height-associated pVLAs, 62.9% of broadly active variants (n = 62) were associated with multiple traits, compared with 51.5% of other pVLAs (n = 530; p = 0.03; **Fig. 5e**). This association supports greater pleiotropic potential among broadly active variants in the height-associated subset.

Having defined endothelial cell diversity within the atlas, we asked whether models trained for individual cell subtypes could reveal variant effects obscured when endothelial cells were analyzed together. Endothelial populations share a vascular identity but occupy distinct physiological microenvironments, including arterial and organ-specific microvascular beds. Subtype-specific fine-tuning resulted in only a modest decline in model performance despite the smaller subtype-specific training sets (**Supplementary Fig. 5b**), indicating that the models retained predictive power while providing greater cellular resolution. We nevertheless hypothesized that divergent subtype regulatory programs would support informative predictions of vascular-bed-specific variant effects.

We fine-tuned seq2PRINT models for eight vascular endothelial subtypes represented by at least 2 million ATAC fragments. Among 1,204 pVLAs with maximal effects in endothelial models, predictions separated from vascular-bed-specific effects (**Fig. 5f**). At the *SMAD6* locus, rs7179372 (which is associated with hip osteoarthritis in a GWAS of hip joint space width and osteoarthritis)^51^, overlapped an endothelial cCRE linked to *SMAD6* and showed its strongest predicted effects in adrenal sinusoidal and sinusoidal endothelial cells, matching subtype-specific accessibility (**Fig. 5g**)^48^. Relative to the broad endothelial model (effect score, −0.38), subtype modeling revealed stronger predicted effects in adrenal sinusoidal (−0.46) and sinusoidal (−0.53) endothelial cells. *SMAD6* regulates Bone morphogenetic protein (BMP) signaling, endothelial junctions and haemodynamic responses^52–55^, making this variant–element–gene relationship a candidate for testing in these specific endothelial cellular contexts.

Despite lower predictive performance at reduced training depth, cell subtype models resolve distinct predicted effects across vascular beds, indicating that regulatory differences remain informative at finer cellular resolution. Subtype-resolved endothelial analysis thus bridges high-resolution cell cartography with variant interpretation, prioritizing precise cellular contexts for functional validation.

## Discussion

The human body functions as an integrated system rather than a collection of isolated organs, requiring a body-wide perspective to fully understand gene regulation. By simultaneously profiling chromatin accessibility and gene expression in nearly half a million nuclei across 21 adult tissues from 4 human donors, we established a cross-tissue atlas to compare general cell programs with tissue-specific adaptations. Across major cell populations, regulatory activity was predominantly driven by core cell identity, while tissue context refined a targeted set of specialized elements and circuits.

Our study maps regulatory architecture across genomic scales, from discrete cis-regulatory elements to broad chromosomal domains. Beyond identifying 161,270 novel candidate cis-regulatory elements absent from ENCODE, and enriched particularly in rare and specialized cell populations, we mapped 871,177 cCRE–gene linked associations. Notably, our atlas extends beyond mapping active elements to delineate constitutively inaccessible domains across primary human cell types. While constitutive heterochromatin is known to remain universally compacted across cellular contexts, and facultative heterochromatin undergoes cell state dependent repression^56^; the distribution and cell-type-specific remodeling of these compartments across human tissues and cells has remained poorly defined. Here, we show that constitutively inaccessible domains cover approximately 10.1% of the analysed genome and overlap 79.3% of sonication-resistant heterochromatin mapped via gradient-seq in fibroblasts, providing orthogonal biochemical validation of their compacted state. Furthermore, these domains are enriched for L1 retrotransposon repeats and recurrent cancer-associated hypomethylation, linking their conserved repressive framework to genome composition and epigenetic dysregulation. Together, these findings expand genome annotation beyond genes and active regulatory elements to encompass broad repressive domains that architecturally structure the non-coding genome.

Non-coding genetic variation underpins much of human disease susceptibility, yet translating statistical associations into functional mechanisms remains a major challenge. Applying seq2PRINT to 548,656 fine-mapped trait-and disease-associated variants, we identified 18,133 high-effect variants predicted to alter chromatin-accessibility (3.3% of those evaluated). Most showed cell-type restricted predicted activity, whereas 6.2% were broadly active across cell lineages. Crucially, 21.4% of high-effect variants fell outside currently annotated cCREs, demonstrating that with higher resolution cell level reference data, sequence-based models can expose novel functional regulatory elements missing from existing reference catalogues. This increased resolution extends even within specialized cell lineages: models trained on vascular endothelial subtypes distinguished predicted variant effects across distinct vascular beds, resolving cellular contexts obscured by broad tissue-level annotations. These findings illustrate how cell-level multi-omic data and predictive modeling can transform genetic associations into mechanistically testable hypotheses - prioritizing specific variants, regulatory elements and target cell populations for base editing and high throughput perturbations. They also motivate expanded sampling and deeper profiling of organs and tissues, developmental stages, physiological exposures, and disease states, where further regulatory effects may emerge.

By establishing a shared molecular coordinate system across tissues and modalities, our study provides a unified reference for integrating current and future single-cell, spatial, and perturbational datasets. Predictive modeling is rapidly reshaping biological discovery, yet the reach of sequence-to-function frameworks - such as AlphaGenome and emerging virtual-cell models - remains fundamentally bound by the breadth and fidelity of their reference training data. Realizing the potential of predictive biology requires empirical, multi-modal measurements that directly link DNA sequence to regulatory state within defined human cell types. Our atlas addresses this critical gap by co-mapping chromatin accessibility and gene expression across human tissues, providing the cellular resolution and regulatory depth required to train, benchmark, and further advance predictive models.

Looking forward, expanding reference maps across diverse ancestries, developmental stages, physiological states and disease contexts will be essential to map dynamic, context-dependent regulatory programs. Ultimately, this study offers more than a catalogue of regulatory elements. It provides an organism-wide perspective on how human gene regulation is conserved across cell types, specialized by local tissue microenvironments, structured by repressive chromatin, and disrupted by non-coding variation. In doing so it establishes a foundational framework for tracing how genetic variation and epigenetic remodeling propagate across biological scales, from primary DNA sequence to chromatin architecture, cellular state, tissue function, and human disease.

## Methods

### Donor and Sample Characteristics

All samples were selected from donors enrolled as part of the GTEx/ENTEX collaborative project^17^. GTEx tissue samples were derived from deceased donors, with study authorization obtained via next-of-kin consent for the collection and banking of de-identified tissue samples for scientific research, previously described^57^. All GTEx samples underwent pathology review as part of GTEx study protocol^58,59^ to confirm tissue origin, content and integrity, and were screened for evidence of any disease to ensure that collected biospecimens were normal for the age of the donor, and non-diseased.

### Tissue preparation, dissociation, and nuclei extraction

Single nuclei were isolated from frozen tissue using a modified lysis buffer to minimize mitochondrial contamination with a detailed protocol^19^. Briefly, approximately 100 mg of frozen tissue was obtained per sample to support multiple extraction protocols, with 20-30 mg aliquots prepared for each extraction. Samples were kept frozen throughout cutting and weighing. Tissues were mechanically dissociated and incubated in an NP-40 based lysis buffer on ice, followed by filtration through a 70 µm strainer and then a 30 µm strainer. The nuclei were then washed and counted using a hemocytometer to ensure a viable and debris-free suspension. A target of 10,000 nuclei per sample was loaded into the Chromium controller to generate single-nuclei GEMs, and multiome profiling was performed using the 10X Genomics Chromium Single Cell Multiome ATAC + Gene Expression kit per the manufacturer’s protocol.

### Library Preparation and Sequencing

Libraries were prepared using the 10X Genomics Chromium Single Cell Multiome ATAC + Gene Expression kit (CG000338). Library quality was assessed on an Agilent Bioanalyzer 2100 with the High Sensitivity DNA Kit (#5067-4626),and concentrations were measured by qPCR using Kapa Library Quantification Kit (#0796014001). Pooled libraries were sequenced on Illumina NovaSeq S2 with paired-end 150 bp reads, targeting at least 50,000 read pairs per nucleus for scATAC-seq and 25,000 read pairs per nucleus for scRNA-seq.

### Multi-omic data processing

Raw snRNA-seq and snATAC-seq reads from the 10x Genomics Multiome platform were demultiplexed with Cell Ranger (v8.0.1)^60^ and Cell Ranger ATAC (v2.1.0)^61^, respectively, and aligned to the GRCh38/hg38 reference genome with GENCODE v39 gene annotation. For snRNA-seq, we applied CellBender (v0.3.2)^62^ to remove ambient RNA and retain high-confidence nuclei, followed by sample- and tissue-specific quality control using Scanpy (v1.11.5)^63^. Low-quality nuclei and putative doublets were iteratively removed based on standard library complexity and mitochondrial/ribosomal content metrics, probabilistic doublet detection, and visual inspection of low-dimensional embeddings. After filtering, 411,952 high-quality nuclei were retained. Counts were normalized and log-transformed, highly variable genes were selected, and batch effects within each tissue were corrected using Harmony (v0.0.9)^64^ before dimensionality reduction and clustering. Cell lineage, cell type and Leiden-derived subcluster annotations were initially assigned using curated marker gene sets with ScType^65^, followed by manual expert refinement.

For snATAC-seq, aligned fragments were first screened with a custom script to remove PCR chimeric reads, and then processed with SnapATAC2 (v2.8.0)^66^ to perform quality control based on fragment counts and transcription start site (TSS) enrichment, and to identify and remove doublets using Scrublet^67^-based scores. After filtering, 234,560 high-quality nuclei were retained.

We then integrated the RNA and ATAC modalities, cross-checking and excluding nuclei flagged as doublets in either modality, resulting in 393,685 RNA cells and 226,859 ATAC cells. GLUE (v0.4.0)^20^ was used to obtain a joint embedding, transfer RNA-derived cell annotations to ATAC cells, and generate a unified UMAP representation. ATAC peaks were initially called with MACS3 (v3.0.4)^68^ (FDR<0.05) across 195 tissue-cell type clusters and resized to 300bp. Clusters containing fewer than 300 cells or fewer than 1 million fragments were excluded from the peak-merging step, leaving 160 clusters. Raw peaks from the retained clusters were merged, and the strongest peak within each merged region was selected as the representative peak only if its best q-value was <10⁻³. This procedure yielded 1,085,062 consensus peaks, which we considered candidate *cis*-regulatory elements (cCREs).

The full details of the processing workflow design are provided in **Supplementary Note 1**. Details of preprocessing, quality-control thresholds, and filtering decisions are provided in **Supplementary Note 2** and **Extended Data Table 1**. Final cluster annotations and key marker genes are summarized in **Extended Data Table 2**. Executable methods are available under **Code Availability**.

### Classification of candidate *Cis*-Regulatory Elements (cCREs)

We classified cCREs by proximity to transcription start sites (TSSs) using GENCODE v39 annotations. For each TSS, we defined a core promoter window of TSS ±200 bp and labeled any 300-bp cCRE overlapping this window as “TSS-overlap”. Among the remaining cCREs, those whose genomic span lay within TSS ±2 kb of the nearest TSS were labeled “TSS-proximal”, and all other cCREs were labeled “TSS-distal”.

### Comparison of cCREs with external datasets

We compared our cCREs with the ENCODE Registry of candidate *Cis*-Regulatory Elements (2,348,854 elements in total)^18^ using BEDTools (v2.31.1; bedtools intersect)^69^ with a ≥1 bp overlap. We similarly intersected our cCREs with experimentally validated human enhancers from the VISTA Enhancer Browser^26^ and with ATAC peaks from a recent single-cell ATAC-seq atlas^5^, and quantified the fraction of elements from each external set overlapping our cCREs. A list of novel cCREs is provided in **Extended Data Table 3**, and the complete list of cCREs is available under **Data Availability**.

### cCRE activity breadth across tissue-cell type combinations

To evaluate the degree of specificity or sharing of cCREs, we overlapped the consensus cCREs with the raw peaks called for each of the 195 tissue-cell type clusters. For each consensus cCRE in each cluster, we assigned a binary label indicating whether the cCRE overlapped at least one raw peak in that tissue-cell cluster. For each cCRE, we then counted the number of tissue-cell type clusters in which it was active and expressed this value as the percentage of tissue-cell type combinations in which the cCRE was active. Analogous activity measures were calculated for AER and ADR domains by determining whether each domain contained at least one active cCRE in a given tissue-cell type combination.

To assess saturation, we calculated the fraction of cCREs that overlapped previously defined ubiquitous representative DNase I hypersensitive sites (rDHSs)^25^ as a function of the number of tissue–cell type combinations considered.

### TF activity and regulatory logic

For each tissue-cell type combination, we quantified transcription factor (TF) activity using chromVAR (GPU version under seq2PRINT package^7^). Starting from the consensus cCRE set, we constructed a binary peak-by-motif matrix using TF motif position weight matrices (PWMs) from CisBP database^70^. This matrix, together with fragment counts for each cCRE, was used to calculate chromVAR deviation scores. chromVAR was run with GC-content and accessibility-bias correction enabled. Matched background cCRE sets were used to control for technical and sequence-composition confounders. The resulting deviation Z-scores provided a TF-specific measure of inferred motif accessibility for each tissue-cell type combination.

To relate inferred TF activity to transcriptional output, following previous analysis^22^, we calculated, for each TF, the correlation between its chromVAR deviation score and expression level across tissue-cell type combinations. These correlations were used to evaluate TF-specific regulatory logic and to identify TFs for which inferred chromatin activity was concordant with transcriptional expression. ChromVAR deviation scores by tissue-cell type combination and TF-expression correlation coefficients are provided in **Extended Data Table 4**.

### CG content of cCREs and AER/ADR domains

To characterize sequence composition of cCREs, we computed mono-CG (C% + G%) and di-CG (CpG%) content for each element. For each cCRE, we extracted the genomic sequence and calculated the fractions of C and G mononucleotides and CpG dinucleotides. These features were then summarized by cCRE group (e.g., TSS, TSS-proximal, TSS-distal, TE-derived vs non-TE-derived) for downstream comparisons.

The same analyses were performed for AER and ADR domains.

### Cross-species alignment breadth and sparsity

To characterize cross-species alignment patterns more directly, we used the 241-species multiple-sequence alignment generated by the Zoonomia Consortium^71^. For each nucleotide position in the human genome, we recorded whether the position aligned to at least one of the other 240 species. For each cCRE and species, we then counted the number of cCRE bases aligned in that species and expressed this value as a fraction of the total cCRE length.

Following the previously described strategy^72^, from these values, we derived two measures for each cCRE. (i) N_1_: the number of species in which ≥90% of bases within the cCRE aligned and (ii) N_2_: the number of species in which ≤10% of bases aligned. These measures were used to classify cCREs according to the breadth and sparsity of their cross-species alignment patterns. cCREs were assigned to one of three predefined groups. G1, a highly conserved group, comprising cCREs with N_1_ ≥ 120 and N_2_ ≤ 25; G2, an actively evolving group, comprising cCREs with 20 ≤ N_1_ ≤ 50 and N_2_ ≤ 120; and G3, a primate-specific group, comprising cCREs with N_1_ ≤ 50 and N_2_ ≥ 180.

### PhyloP and phastCons conservation profiles

We used a list of phyloP and phastCons conservation scores to assess the evolution conservation of regulatory elements. The conservation tracks included 234-way primate phyloP^71^, 241-way placental mammal phyloP^71^, 100-way vertebrate phyloP^73^, and 100-way vertebrate phastCons scores^73^.

For each cCRE, we extracted phyloP scores in a 4-kb window centered on the cCRE midpoint (±2kb), binned the region into 10-bp bins, and averaged scores within each bin. We then generated aggregate conservation metaprofiles by averaging these scores across cCREs in predefined groups, including TSS, TSS-proximal and TSS-distal cCREs; cCREs overlapping versus not overlapping ENCODE cCREs; and transposable-element-derived versus non-transposable-element-derived cCREs.

To assess the robustness of conservation patterns across phylogenetic comparisons and conservation metrics, we repeated these analyses using each of the four conservation tracks described above. Because primate-specific cCREs may have limited representation in the placental-mammal and vertebrate alignments, and may therefore have reduced power for conservation analysis, we repeated the metaprofile analyses after restricting the cCRE set to the G1 highly conserved group.

### Identification and cell-type enrichment of accelerating cCREs

We classified cCREs according to their cross-species alignment breadth and sparsity, as described above. We restricted subsequent analyses to TSS-distal cCREs assigned to the G3 group, which was previously defined as the primate-specific group. These cCREs were referred to as accelerating cCREs for downstream analyses.

To assess cell-type enrichment, we quantified the proportion of ATAC-seq fragments overlapping G3 TSS-distal cCREs for each cell type. Specifically, for each cell type, we calculated the fraction of all fragments overlapping the selected accelerating cCREs, using the corresponding total number of fragments as the denominator. Cell types with a higher proportion of fragments mapping to these cCREs were considered enriched for accelerating cCRE activity.

### Identification of marker cCREs

For dynamic analyses, we identified marker cCREs at the level of tissue–cell type pairs. For each pair and each cCRE, we calculated the difference between the mean normalized ATAC–seq signal in that pair and the mean signal across all other tissue–cell type pairs. cCREs were ranked according to this contrast, and the 500 highest-ranked cCREs for each pair were retained as pair-specific marker cCREs. The union of these marker sets comprised 42,619 cCREs and was used for downstream analyses. Normalized ATAC–seq signal across this union was visualized for all tissue–cell type pairs.

For selected cell types identified across multiple tissues, we additionally identified marker genes and visualized ATAC-seq signal at their associated cCREs across cells of the corresponding cell type, stratified by tissue of origin.

### Classification of cell type-dominant and tissue-dominant cCRE activity

To evaluate specificity of regulatory activity, we focused on five major cell types (blood vascular endothelial cell, macrophage, T cell, fibroblast, and smooth muscle cell) that were robustly identified across most tissues (more than 10 tissues). For each cell type, we aggregated active cCREs across tissues and calculated, for each cCRE, the number of tissues in which it was active.

A cCRE was classified as cell type-dominant for a given cell type if it was active in at least half of the tissues containing that cell type and showed substantially lower activity in other cell types within the tissue in which its activity was highest. cCREs that were active in exactly one tissue for that cell type were classified as tissue-dominant.

### cCRE-gene link analysis

To infer cCRE-gene links, we developed fastDORC, an accelerated implementation of DORC (Domains of Regulatory Chromatin) framework^29^ (**Supplementary Note 3**) and applied it to our joint snRNA-seq and snATAC-seq profiles (**Supplementary Note 4**). For each gene, we considered all cCREs within ±250 kb of its TSS and quantified *cis*-associations between gene expression and chromatin accessibility across matched cells using the DORC pipeline.

cCREs showing significant positive or negative associations with a gene were retained as linked regulatory elements using an FDR threshold of 0.05. For each gene, the DORC score was calculated as the summed accessibility across its linked cCREs. cCRE–gene linking and DORC calling were performed independently within each cell type. Links were subsequently compared across cell types to assess their reproducibility and sharing.

The complete set of DORC cCRE–gene links and the genes identified as DORC genes is provided under **Data Availability**.

### Transposable element annotation

Transposable element (TE) annotations were generated by running RepeatMasker (v4.0.7)^74^ on the human reference genome using Repbase (v30.10)^75^ consensus sequences as the TE library, with parameters “-s -no_is-nolow -norna -pa 32 -e ncbi -cutoff 255 -div 40 -frag 20000”. This produced both genomic TE annotations and alignments of individual TE insertions to their corresponding consensus sequences.

### Classification of TE-derived cCREs

Genomic TE annotations were used to classify regions as TE-derived or non-TE-derived by intersecting the center of cCRE with TE annotations using BEDTools (bedtools intersect)^69^, requiring at least 1 bp overlap. A cCRE was classified as TE-derived if its midpoint overlapped a TE annotation by at least 1 bp; all other cCREs were classified as non-TE-derived. The same procedure was applied to AER and ADR domains, where applicable.

The RepeatMasker alignment output was used to map TE-derived cCREs to their corresponding TE consensus sequences, enabling analysis of the positional distribution of regulatory elements along consensus coordinates.

### TE-family enrichment

To quantify enrichment of individual TE families, we compared their representation among cCREs and AER/ADR domains with their representation in the corresponding genomic background. Enrichment was calculated as the ratio of the observed proportion of elements assigned to a given TE family to its expected proportion in the background set.

### Definition of ATAC-depleted regions (ADRs)

Motivated by prior evidence on large heterochromatic domains^34^, we partitioned the human genome into non-overlapping 100-kb windows. We calculated 50-mer mappability scores^76^ for each window and excluded windows with score <0.5, as well as windows overlapping previously defined problematic regions or the pseudoautosomal regions (PARs). This filtering yielded 26,502 high-quality 100-kb windows.

For each tissue-cell type combination, we quantified ATAC–seq accessibility in each window as the total number of fragments overlapping the window and normalized these values to library size as counts per million (CPM). CPM values were then converted to Z-scores across tissue-cell type combinations and used for unsupervised clustering of windows with similar accessibility profiles.

We next evaluated mean accessibility and cCRE activity across the resulting window clusters. Windows exhibiting high accessibility across many samples and containing active cCREs in most tissue–cell type combinations were classified as ATAC-enriched regions (AERs; n = 3,470). Windows with consistently low accessibility and few or no active cCREs were classified as ubiquitously ATAC-depleted regions (ubi-ADRs; n = 2,690). All remaining windows were classified as cell type-specific ATAC-depleted regions (cts-ADRs; n = 20,342). These cts-ADRs were further subclustered and annotated according to the cell types in which they showed the highest accessibility. The resulting AER and ADR annotations are provided in **Extended Data Table 5**.

### Comparison with sonication-resistant heterochromatin

We compared constitutively inaccessible domains with sonication-resistant heterochromatin domains previously mapped in fibroblasts by Becker *et al.*^32^ A constitutively inaccessible domain was considered overlapping a reference interval if the two regions shared at least 1 bp.

### DNA methylation changes in cancer

We downloaded Illumina DNA methylation array data from the TCGA data portal^77^ and restricted analyses to TCGA projects with both tumor and normal samples, resulting in 9,051 samples across 25 projects. DNA methylation β-values were generated from raw IDAT files for tumor and normal samples across these cancer projects using methylprep (v1.7.1)^78^.

For each sample, we calculated two measures. (i) the mean β-value within each individual chromatin-domain region and (ii) a whole-array global mean β-value averaged over all probes on that sample. To account for differences in global methylation levels between cancer types and samples, we normalized the mean β-value of each region by subtracting the corresponding sample-specific whole-array mean β-value.

Within each project, we then computed, for every individual domain region, the difference between the mean global-mean-normalized β of tumor samples and that of normal samples (requiring ≥2 tumor and ≥2 normal samples with data for that region). Regions were included only when methylation data were available for at least two tumour and two normal samples. Finally, we averaged these region-level tumour–normal differences across all regions belonging to each domain subgroup. Thus, each genomic region contributed equally to the subgroup-level estimate, irrespective of the number of methylation probes it contained, yielding a project-by-domain-subgroup delta-methylation matrix.

### Sequence-based modeling of chromatin accessibility

We used seq2PRINT^7^ to model sequence-encoded chromatin accessibility across cell types. For each cell type, we aggregated ATAC-seq fragments and called pseudobulk peaks. These peaks were used as training targets and are referred to here as cell-type-specific regulatory regions. Following the recommended seq2PRINT framework, we extracted 4,096-bp genomic windows centered on each peak and used the corresponding DNA sequences as model inputs. Pseudobulk ATAC-seq accessibility at the central peak served as the prediction target.

Models were trained for 35 cell types with at least 10 million ATAC-seq fragments. This threshold was selected to provide sufficient coverage at individual peaks and stable pseudobulk accessibility profiles. Genomic regions were partitioned into training, validation and test sets, with held-out genomic regions used to reduce overfitting and assess generalization to unseen loci. Model performance was evaluated using Pearson correlations between observed and predicted accessibility signals and between observed and predicted transcription-factor (TF) footprint profiles (**Supplementary Fig. 5a**).

For blood vascular endothelial cells, we used the corresponding cell-type model as a shared backbone and fine-tuned only the final task-specific layers for eight subtypes with at least 2 million ATAC-seq fragments. This approach enabled subtype-specific refinement while preserving a common sequence-processing backbone across related subtypes. Fine-tuned models were evaluated on held-out genomic regions by comparing predicted accessibility and TF footprint profiles with the corresponding pseudobulk ATAC-seq signals using Pearson correlation. Performance metrics for all fine-tuned models are shown in **Supplementary Fig. 5b**.

### Validation against independent chromatin-accessibility QTLs

To evaluate the accuracy and generalizability of sequence-based variant-effect predictions, we trained seq2PRINT models using bulk ATAC-seq data from Corces et al.^45^, as well as ENCODE chromatin-accessibility data from monocytes (ENCSR009WQK, ENCSR410OWB, pooled into these two snATAC-seq into one pseudobulk), B cells (ENCSR903WVU, ENCSR610AQP), CD8 T cells (ENCSR283LPH, ENCSR392YGP), and CD4 T cells (ENCSR452COS, ENCSR841LHT)^35^. For ENCODE B, CD4 T and CD8 T cells, we trained models independently on two datasets representing the same cell type. The predicted allelic effects from the two independently trained models were averaged for each variant-peak pair to obtain a single cell-population-specific prediction.

We compared these predictions with independently measured chromatin-accessibility quantitative trait locus (caQTL) effects from the CIMA atlas of 428 individuals^15^. We quantified agreement using Pearson and Spearman correlation within each matched cell population. For each variant and matched cell population, the predicted allelic effect was calculated as the difference between the model-predicted accessibility for the alternative and reference alleles. Observed caQTL effects were represented by the reported median allelic slope. Variants were matched between the seq2PRINT predictions and the CIMA caQTL dataset using genomic position and alleles, and effect directions were harmonized to the same reference and alternative alleles. Agreement between predicted and observed effects was assessed separately for B cells, CD4 T cells, CD8 T cells and monocytes using Pearson and Spearman correlation coefficients. Only variants with both a predicted effect and an available CIMA caQTL estimate were included. We estimated 95% confidence intervals by bootstrapping over clusters of SNP-peak pairs (n = 10,000 resamples with replacement), taking the 2.5th and 97.5th percentiles. Clusters were defined by single linkage of pairs that shared a variant or peak, or whose variants lay within 100 kb.

To assess the dependence of variant-effect prediction on training-data depth, we used the Corces et al.^45^ data and downsampled the training fragments to nested subsets of 2, 5, 10, 25, 35 million unique fragments. Models were retrained at each sequencing depth using the same architecture and genomic training, validation and test partitions as the full-depth models. Predicted allelic effects and TF footprint profiles were generated at each depth. Performance was evaluated by comparing predicted accessibility effects with CIMA caQTL effects using Pearson and Spearman correlation coefficients, and by comparing full-depth observed and predicted chromatin-accessibility and TF footprint profiles. These results were compared with those from models trained using the maximum available sequencing depth for each cell population: 34.9 million unique fragments for monocytes and 46.4 million unique fragments forCD4 T cells.

### *In silico* prediction and enrichment of GWAS variant effects

We quantified cell type-specific regulatory effects of GWAS fine-mapped variants using credible sets from Open Targets^46^. We downloaded credible sets comprising 588,618 variants across 48,828 studies and restricted analyses to studies with ≥50 SNPs having PIP >0.1, yielding 6,631 studies. Of these variants, 548,656 were represented in the seq2PRINT input space and were analyzed further.

For each eligible SNP, we performed in silico mutagenesis with each trained seq2PRINT model and calculated a predicted cell-type-specific effect score by comparing the model output for the reference and alternative alleles. This produced one predicted effect score (Δ) for each SNP-cell type pair.

To test enrichment of predicted regulatory effects, we evaluated a grid of PIP thresholds (0.1-1.0). For each study, cell type, and PIP threshold, we computed the mean absolute Δ across SNPs with PIP ≥ threshold (requiring ≥50 SNPs). This value was compared with a leave-one-study-out (LOSO) background constructed from credible-set variants in all other studies at the same PIP threshold and for the same cell type. The comparison yielded an analytic Z-score and a two-sided *p-value* based on the standard normal distribution.

For each study, we selected an optimal PIP threshold using the median-Z procedure described in **Supplementary Note 5** and **Supplementary Fig. 6**. Briefly, the median Z-score across cell types was used only for threshold selection. All reported effect sizes and *p-values* were then recalculated at the selected threshold using an independent LOSO background. Multiple testing across the complete study-by-cell type grid was controlled using the Benjamini-Hochberg procedure applied to the final set of Z-derived *p-values*. To account for selecting the PIP threshold that maximized the cross-cell-type statistic, we will permute PIP values within each study, repeat the complete threshold-selection procedure and compare each observed optimized statistic with its empirical optimized null distribution. The completed analysis will report the number of permutations, empirical *P-value* resolution and concordance with the analytic results.

The study-by-cell type Z-scores calculated at the selected PIP thresholds were used as the primary measure of cell-type-specific enrichment of predicted variant effects. Overall, 2,170 studies were significantly associated with at least one of the 35 modeled cell types. Study–cell type associations and corresponding Z-scores are reported in **Extended Data Table 6**.

### Identification of predicted variants with large accessibility effects

For each of the 548,656 variants represented in the seq2PRINT input space, we obtained in silico allelic effect scores (Δ) from each of the 35 trained cell-type-specific models. The delta score was defined as the difference between the predicted chromatin-accessibility signal for the alternative and reference alleles.

To summarize the maximum predicted regulatory effect of each variant across cell types, we calculated the maximum delta score across all 35 cell types. Variants were ranked according to maximum delta score. We classified variants with maximum delta score≥ 0.2 as predicted variants with large accessibility effects (pVLAs), yielding 18,133 pVLAs. The cell type associated with the largest absolute effect score was assigned as the variant’s putative primary cell type. For visualization, variants were represented in the joint cell-type embedding according to this cell-type assignment and colored by maximum delta score.

To quantify the cell-type specificity of predicted effects, we calculated a Gini index from the absolute delta scores across the 35 modeled cell types for each variant. Higher Gini values indicate that the predicted effect is concentrated in a smaller number of cell types, whereas lower values indicate a more broadly distributed effect across cell types. pVLAs with a gini index of delta score < 0.4 are classified as ubiquitous pVLAs (u-pVLAs).

For analyses focused on blood vascular endothelial cells, we additionally identified the subset of pVLAs for which the maximum absolute delta score was observed in one of the endothelial-cell models. This yielded 1,204 endothelial-cell-associated pVLAs. These variants were further evaluated across endothelial subtypes using the subtype-specific seq2PRINT models.

### Enrichment of pVLAs across Open Targets studies

To identify phenotypic studies enriched for pVLAs, we analyzed credible-set variants from all eligible Open Targets studies. For each study, we used the study-specific optimized PIP threshold (**Supplementary Note 5**) determined in the preceding GWAS variant-effect analysis. For each study, variants with PIP values at or above its optimized threshold were retained. Studies were included only when at least 50 variants satisfied this threshold.

We tested whether pVLAs were over-represented among the retained variants for each study using a one-sided Fisher’s exact test. The background universe was restricted to variants in the seq2PRINT input space that were eligible for pVLA classification and evaluated using the corresponding study-specific PIP threshold. For each study, we constructed a 2 × 2 contingency table comparing pVLA and non-pVLA status among variants included in the study and the background set.

For each study, we calculated the enrichment odds ratio and the corresponding Fisher’s exact-test *p-value*. An odds ratio greater than 1 indicated enrichment of pVLAs among the study-associated variants.

## Data Availability

The raw data in fastq format and processed 10x multi-omic datasets generated in this study will be made available through the Chan Zuckerberg Initiative (CZI) CELLxGENE database upon publication (https://cellxgene.cziscience.com). The datasets will also be available on the GTEx portal (www.gtexportal.org), along with additional data tables from this study. Expanded phenotype metadata for the donors in this study will be available from the GTEx study, through the database of Genotypes and Phenotypes (dbGaP) (https://dbgap.ncbi.nlm.nih.gov/) under the study accession number phs00424.v11.p2.

Additional data supporting the findings of this study are available from the corresponding author upon reasonable request.

## Code Availability

The tissue-aware preprocessing and annotation workflow, GLUE-based label-transfer implementation will be available at https://github.com/KailiBio/Multiomic-workflow. Custom scripts for downstream analyses will be available at https://github.com/KailiBio/human-adult-cross-tissue-scmultiome-analysis.

Any additional resources or materials generated in this study are also available from the corresponding author upon reasonable request.

## Acknowledgements

We thank the generosity of the donor families for providing such generous gifts and making this study possible. We also thank Kenneth Zaret for helpful discussions and insights regarding the use of ATAC-seq data to identify heterochromatin regions. We thank Jared Nedzel and Joseph Okonda for their assistance with data sharing and visualization through the GTEx Portal. The results presented here are based in part on data generated by The Cancer Genome Atlas (TCGA) Research Network. https://www.cancer.gov/tcga. Schematic illustrations were created with BioRender.com.

## Author Contributions

K.G.A., F.A., A.G.A., and A.S. conceived the project and designed the experiments. A.G.A., A.S., and M.K.T. isolated nuclei from frozen tissue and generated sequencing libraries. K.F. led the data analysis, with contributions from N.D., L.D., T.Y., Z.C., and R.Z.. N.D. performed primary sequencing-data processing using Cell Ranger, conducted ScrinVex analyses, and, together with K.F., performed RNA and ATAC quality-control assessments. L.D. performed CellBender-based background correction. Z.C. developed and integrated a cell-embedding-based cell-annotation function into GLUE. R.Z. developed an updated implementation of fastDORC and contributed to the seq2PRINT application. R.Z., X.Z. and Z.C. performed VEP validation of seq2PRINT using the CIMA dataset. T.Y. generated the transposable-element annotation file and contributed methodological expertise to the design and interpretation of the transposable-element analyses. K.F. and J.D.B. wrote the manuscript, with input from all authors. J.D.B., and K.G.A. supervised all aspects of the study.

## Ethical Considerations

All experimental procedures involving human or animal samples were approved by [Institutional Review Board (IRB)/Ethics Committee]. Donor authorization for research was obtained via next-of-kin consent.

---

**Extended Data Figure 1.**
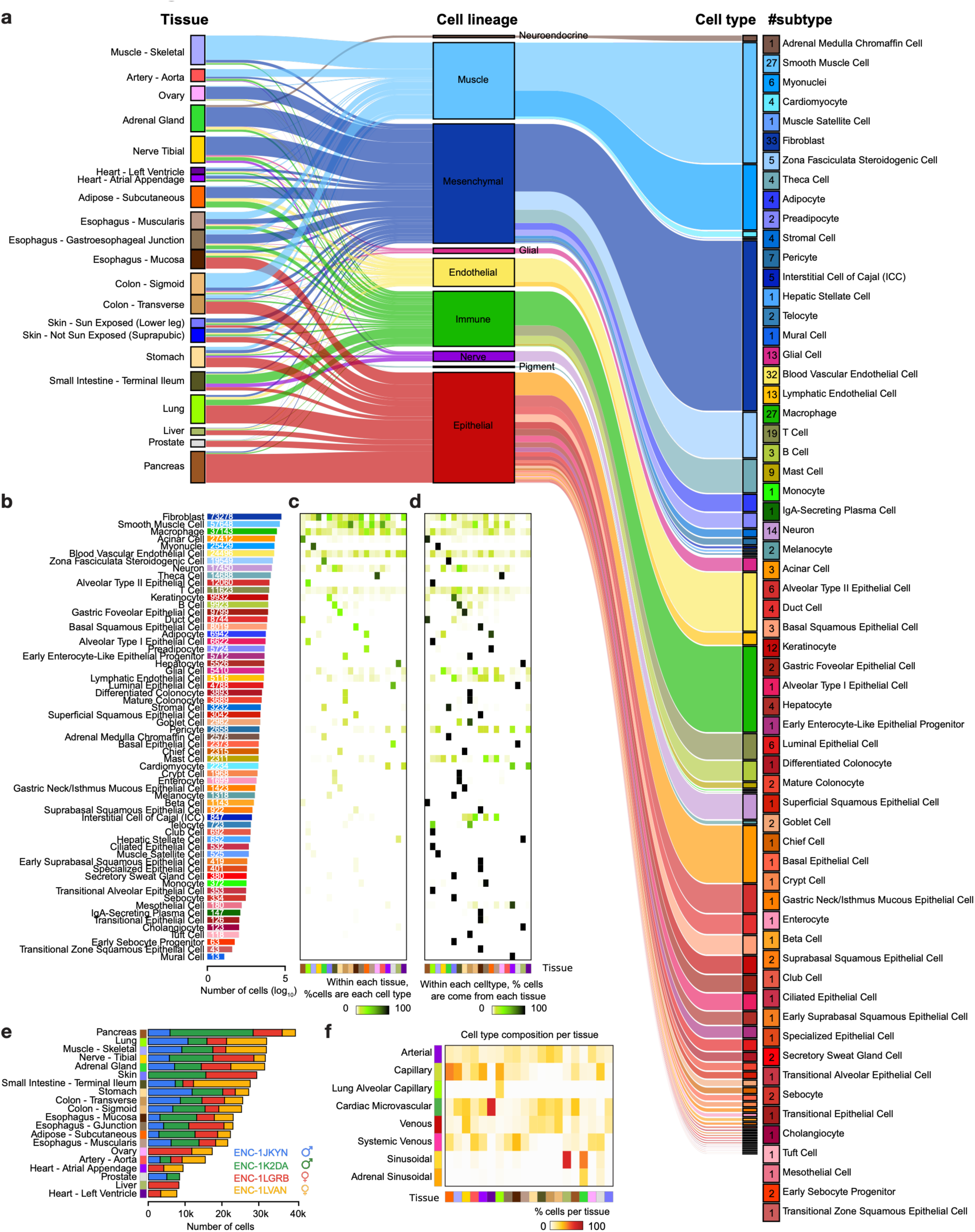
Tissue and donor contributions to broad cell types. **a**. Alluvial plot showing the relationships among tissues, cell lineages, broad cell types and number of subtypes per celltype. **b**. Number of nuclei assigned to each broad cell type, shown on a log_10_ scale. **c**. Heatmap showing, within each tissue, the percentage of nuclei assigned to each broad cell type. **d**. Heatmap showing, within each broad cell type, the percentage of nuclei contributed by each tissue. **e**. Number of nuclei profiled in each tissue, shown as stacked bars colored by donor. **f**. Endothelial cell-type composition of each tissue. Colors indicate the percentage of endothelial nuclei in each tissue assigned to each blood vascular endothelial cell subtype. The color bar below indicates tissue identity.

**Extended Data Fig. 2.**
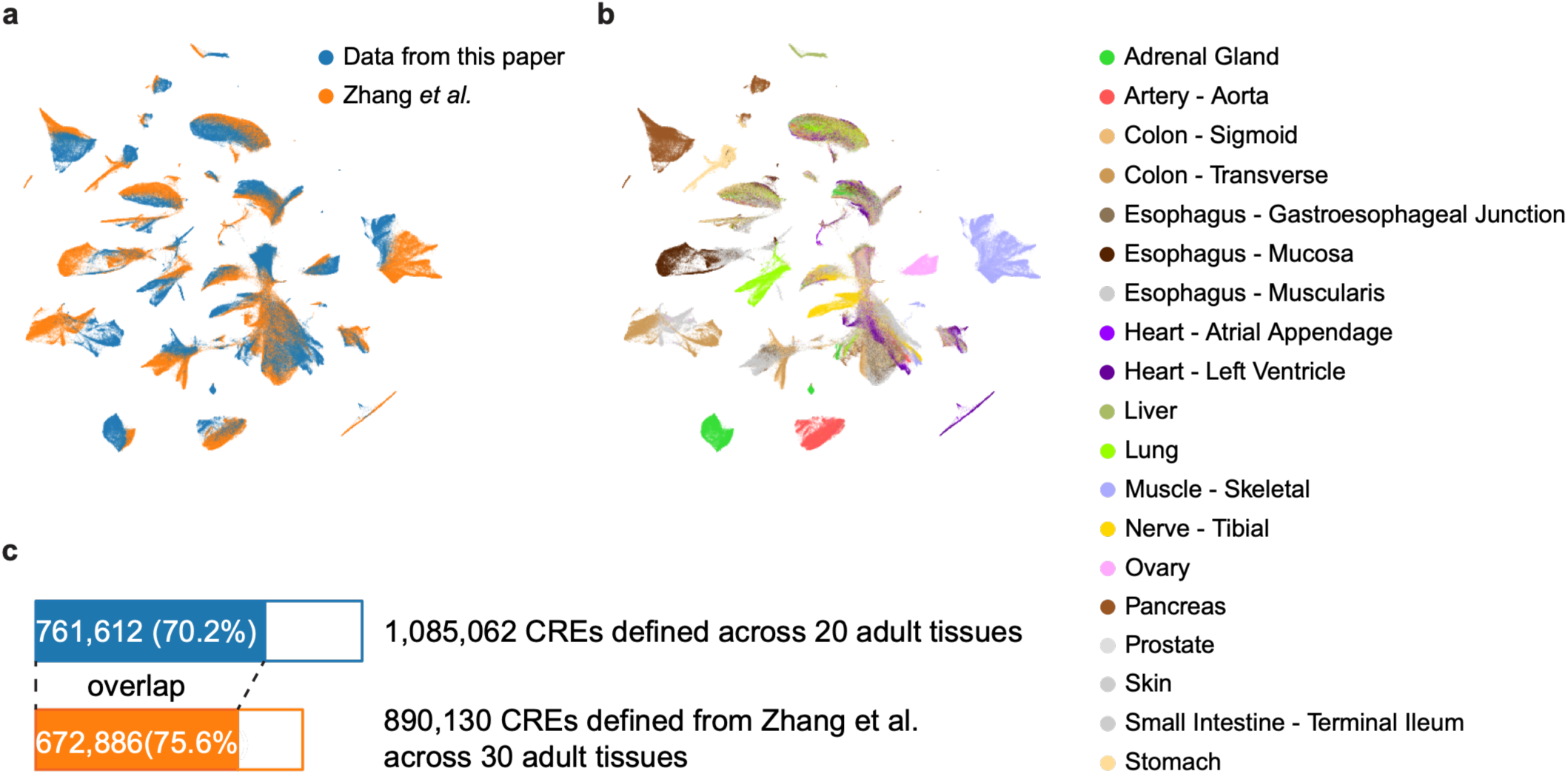
Comparison of candidate *cis*-regulatory elements with an independent single-cell ATAC-seq atlas. **a.** Joint UMAP embedding of nuclei from this study and from the sci-ATAC-seq atlas of Zhang *et al.*, colored by dataset. **b.** The same joint UMAP embedding as panel a, colored by tissue of origin. **c.** Overlap between candidate *cis*-regulatory elements (cCREs) identified in this study and those reported by Zhang *et al*.

**Extended Data Fig. 3.**
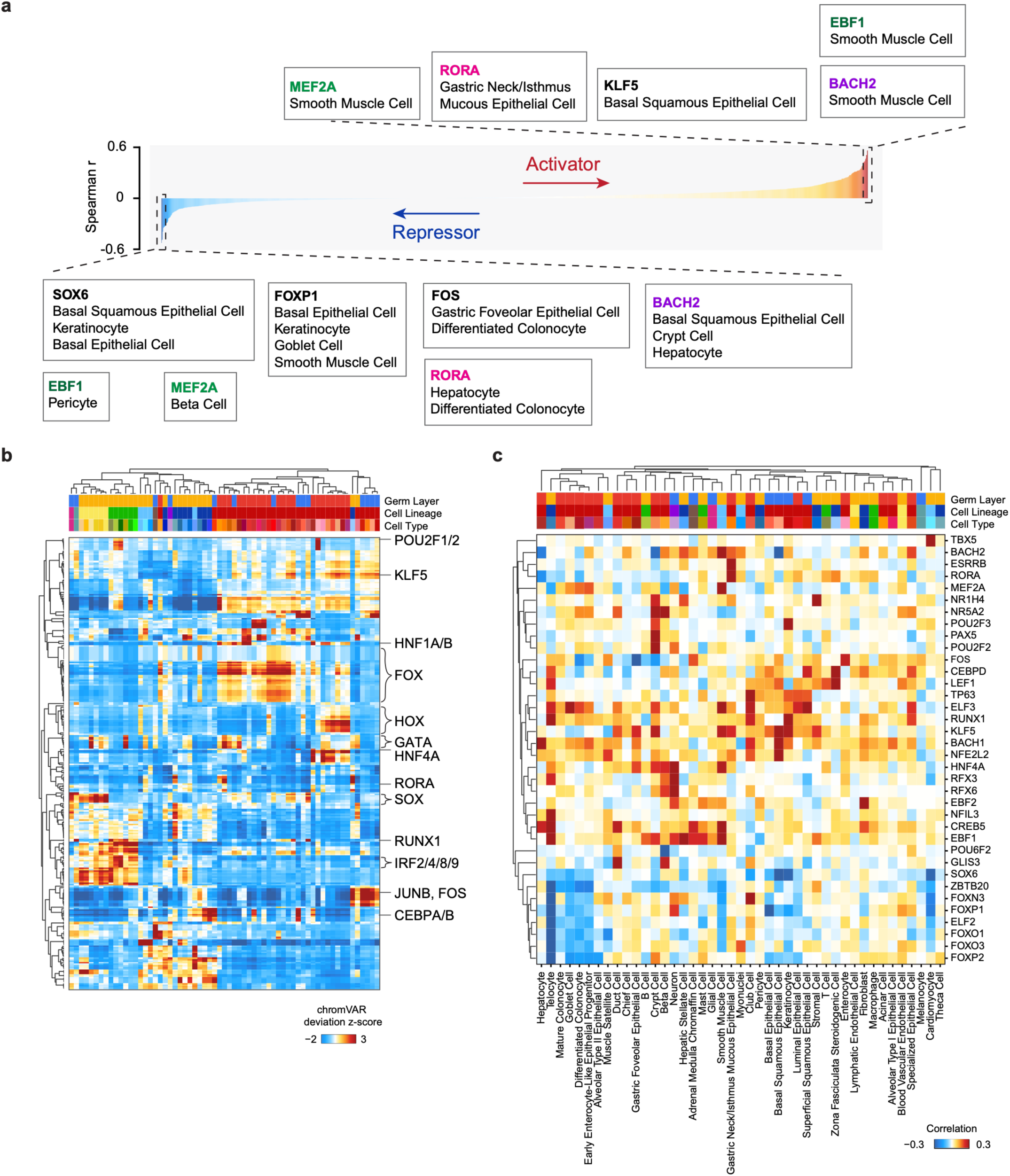
Transcription factor regulatory logic across human cell types. **a.** Spectrum of transcription factor regulatory logic across selected TF-cell-type pairs. TF-cell-type pairs are arranged according to the Spearman correlation between TF expression and chromVAR motif activity, ranging from repressor-like (negative correlation, blue) to activator-like (positive correlation, red). Representative examples of context-dependent regulatory behaviour are indicated, including SOX6, FOXP1, FOS, BACH2, RORA, MEF2A and EBF1. **b**. Heatmap of chromVAR motif-deviation Z-scores across primary human cell types. Rows represent TF motifs and columns represent cell types ordered by hierarchical clustering. Annotation bars indicate germ layer, cell lineage and cell type. **c**. Heatmap of Spearman correlations between TF RNA expression and chromVAR motif-deviation scores across primary human cell types. Positive correlations are shown in red and negative correlations in blue. Rows and columns are ordered by hierarchical clustering, with annotation bars indicating germ layer, cell lineage and cell type. The matrix reveals lineage-associated and cell-type-dependent patterns of TF expression-motif activity coupling.

**Extended Data Fig. 4.**
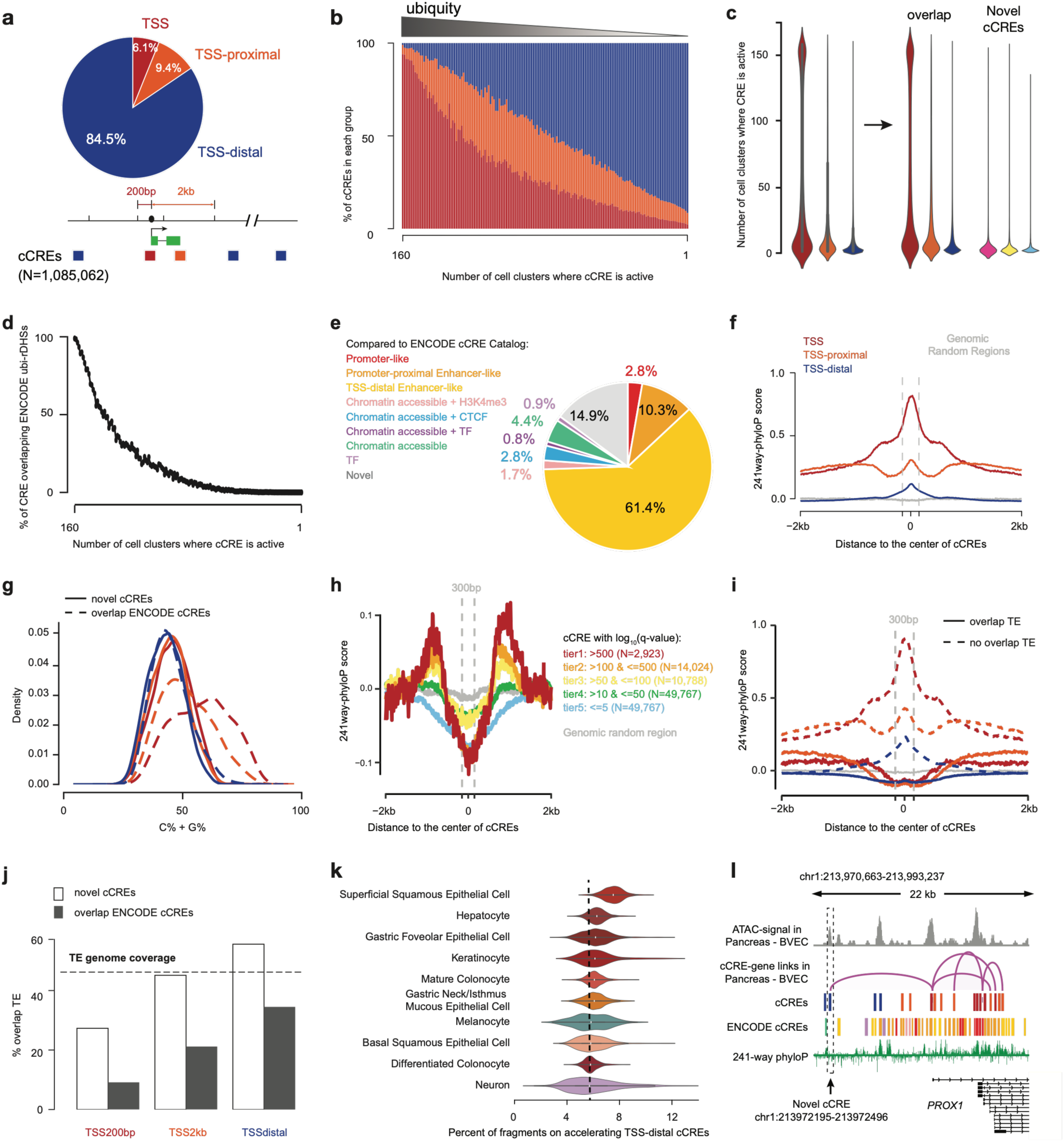
Additional characterization of cCRE activity and regulatory recurrence. **a.** Genomic distribution of 1,085,062 candidate *cis*-regulatory elements (cCREs) relative to GENCODE transcription start sites (TSSs). cCREs were classified as TSS-associated (within ±200 bp of a TSS), TSS-proximal (200 bp-2 kb) or TSS-distal (>2 kb). **b.** Composition of cCREs according to the number of cell clusters in which they are active. Colors indicate TSS-associated, TSS-proximal and TSS-distal cCREs. **c**. Distribution of the number of cell clusters in which individual cCREs are active, stratified by genomic-context group (left), and further stratified by whether overlapping ENCODE cCREs on the right. **d.** Percentage of cCREs overlapping ubiquitously accessible regulatory elements (ubi-rDHSs) reported by Fan *et al*., according to the number of cell clusters in which each cCRE is active. **e**. Classification of cCREs according to overlap with the ENCODE cCRE catalogue. Segments indicate ENCODE cCRE classes and novel cCREs. **f**. Aggregation plots of 241-way placental mammal phyloP scores centred on cCREs, stratified by tiers of peak accessibility significance. The grey line indicates genomic random regions. **g.** Density distributions of monoCG (C%+G%) for cCREs, stratified by genomic-context group and by overlap with the ENCODE cCRE catalogue. Solid lines indicate novel cCREs and dashed lines indicate ENCODE-overlapping cCREs. **h**. Aggregation plot of 241-way placental mammal phyloP scores centred on TSS-distal novel cCREs, stratified by peak log_10_(q-value) tiers. The grey line indicates genomic random regions. **i**. Aggregation plot of 241-way placental mammal phyloP scores centred on cCREs, stratified by overlap with annotated transposable elements. **j.** Percentage of cCREs overlapping annotated transposable elements, stratified by genomic-context group and cCRE novelty. The dashed line indicates the genome-wide fraction of bases annotated as transposable elements. **k**. Percentage of ATAC-seq fragments overlapping TSS-distal, evolutionarily accelerated cCREs in representative cell types. Cell types are ordered according to the proportion of fragments overlapping these cCREs. **l**. Genome-browser view of a novel TSS-distal cCRE at chr1:213,972,195-213,972,496 in pancreatic blood vascular endothelial cells. The cCRE shows cell-type-specific accessibility, is absent from the ENCODE cCRE catalogue and is linked to PROX1 by peak-gene associations.

**Extended Data Fig. 5.**
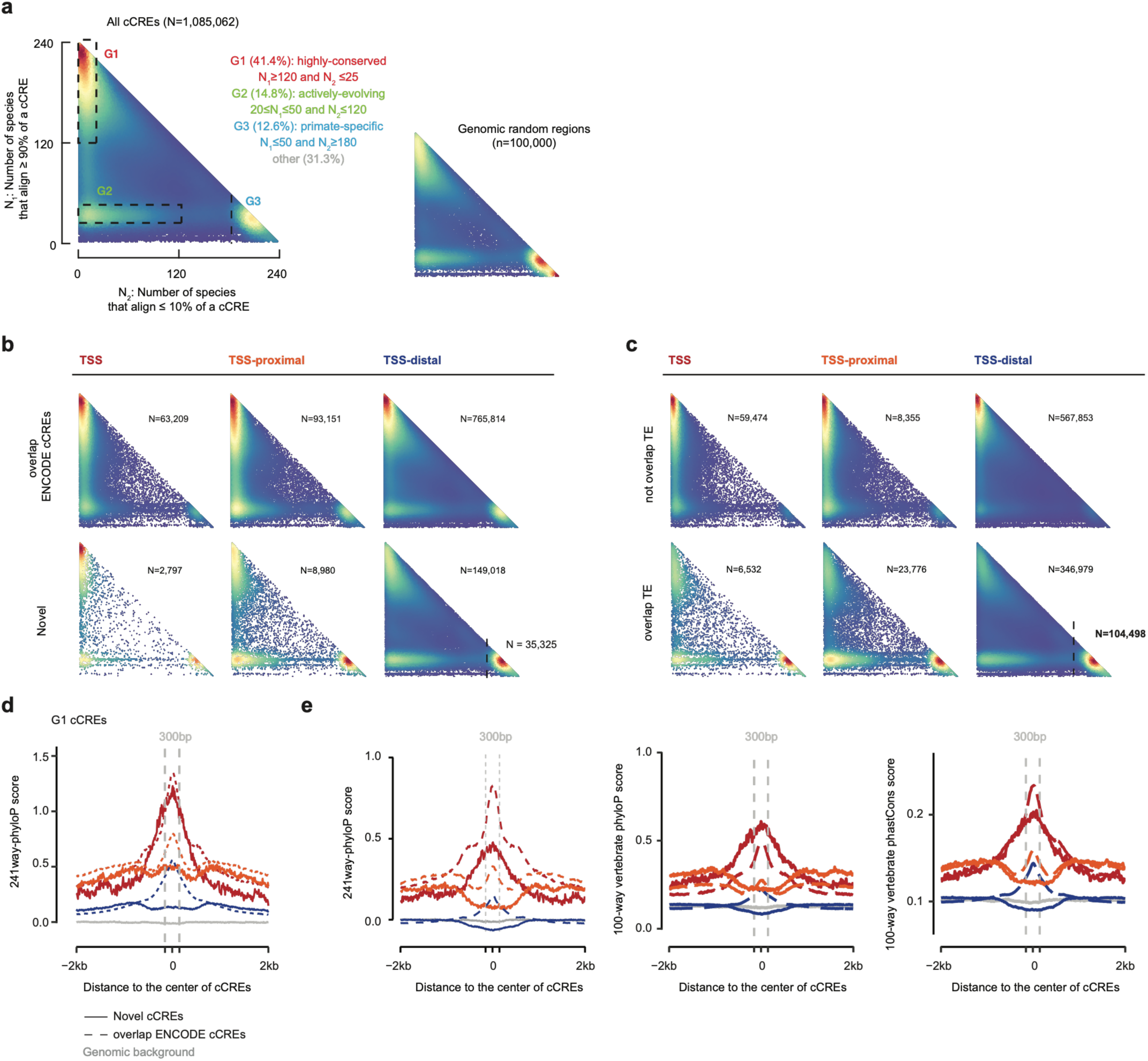
Evolutionary conservation and cross-species alignability of cCREs. **a.** Two-dimensional histograms of cross-species alignability for all cCREs (left) and genomic random regions (right). N₂ denotes the number of species in which ≤10% of cCRE nucleotides align, and N₁ denotes the number of species in which ≥90% of nucleotides align. Dashed regions define highly conserved (G1), actively evolving (G2) and primate-specific (G3) cCREs. **b**. Cross-species alignability distributions for cCREs stratified by genomic-context group and novelty. **c**. Cross-species alignability distributions for cCREs stratified by genomic-context group and overlap with annotated transposable elements. **d**. Aggregation plots of 100-way mammal phyloP centred on G1 cCREs (defined from panel a), stratified by genomic-context group and novelty. Dashed grey lines indicate genomic random regions. **e**. Aggregation plots of 100-way mammal phyloP, 100-way vertebrate phyloP and 100-way vertebrate phastCons scores centred on cCREs, stratified by genomic-context group and novelty. Dashed grey lines indicate genomic random regions.

**Extended Data Fig. 6.**
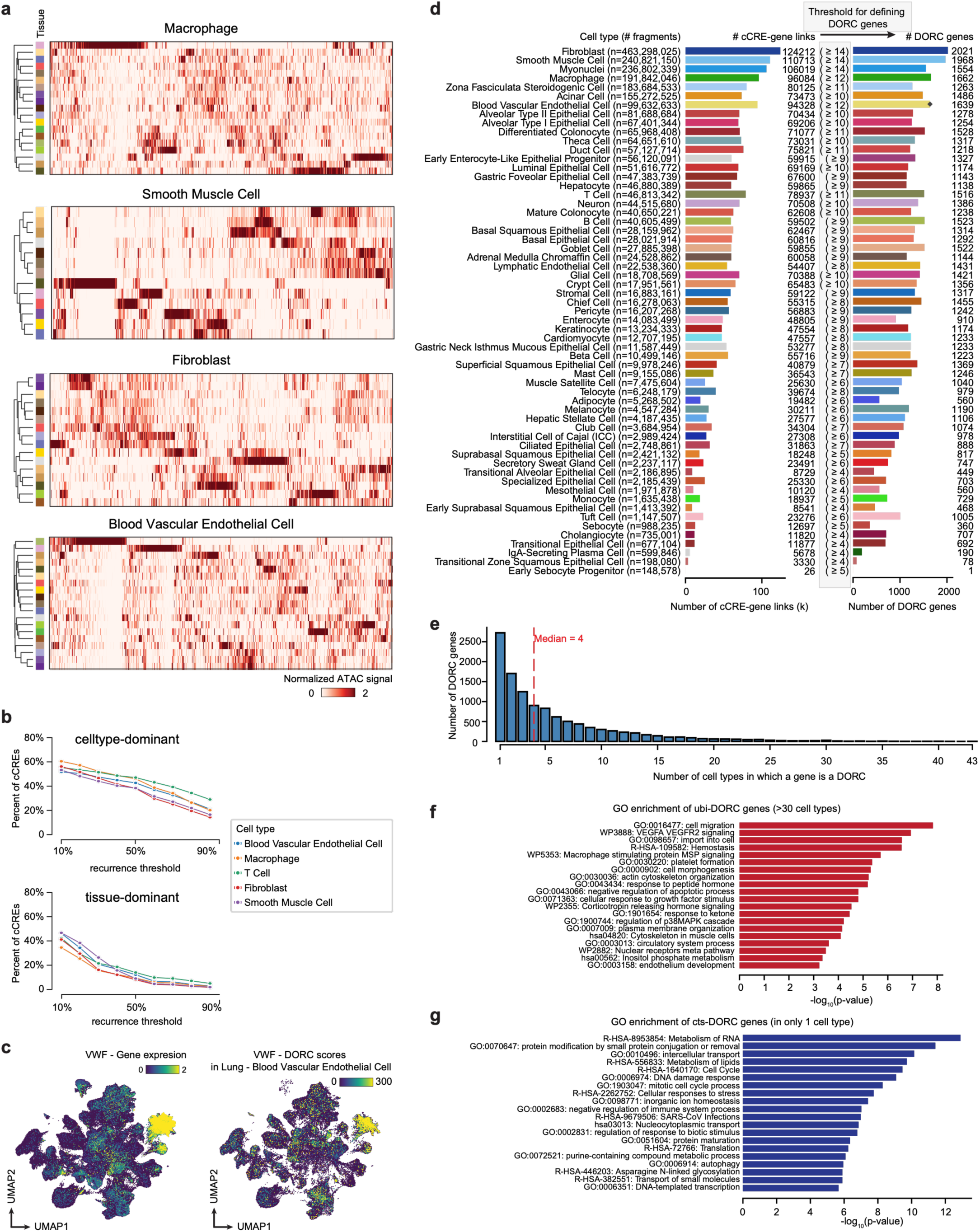
Cell-type-specific cCRE activity and DORC recurrence. **a**. Heatmaps showing normalized ATAC-seq signal for the 500 most variable cCREs in macrophages, smooth muscle cells, fibroblasts and blood vascular endothelial cells. Cell clusters are ordered by hierarchical clustering; annotation bars indicate tissue of origin. **b**. Fraction of peaks classified as cell-type-dominant, tissue-dominant or interaction-dominant as a function of the recurrence threshold. Curves are shown for blood vascular endothelial cells, macrophages, T cells, fibroblasts and smooth muscle cells. **c**. UMAP embeddings of lung vascular endothelial cells colored by VWF expression (left) and DORC scores for VWF (right). **d**. Number of cCRE–gene links (left) and DORC genes (right) for each cell type. Cell types are ordered by the number of ATAC-seq fragments; values in parentheses indicate the minimum number of cCRE–gene links required to define a DORC gene. **e**. Distribution of DORC genes according to the number of cell types in which they were identified. The dashed line indicates the median of 4. **f**. Gene Ontology (GO) terms enriched among ubiquitous DORC genes, defined as genes identified as DORCs in more than 30 cell types. **g**. GO terms enriched among cell-type-specific DORC genes, defined as genes identified as DORCs in a single cell type.

**Extended Data Fig. 7.**
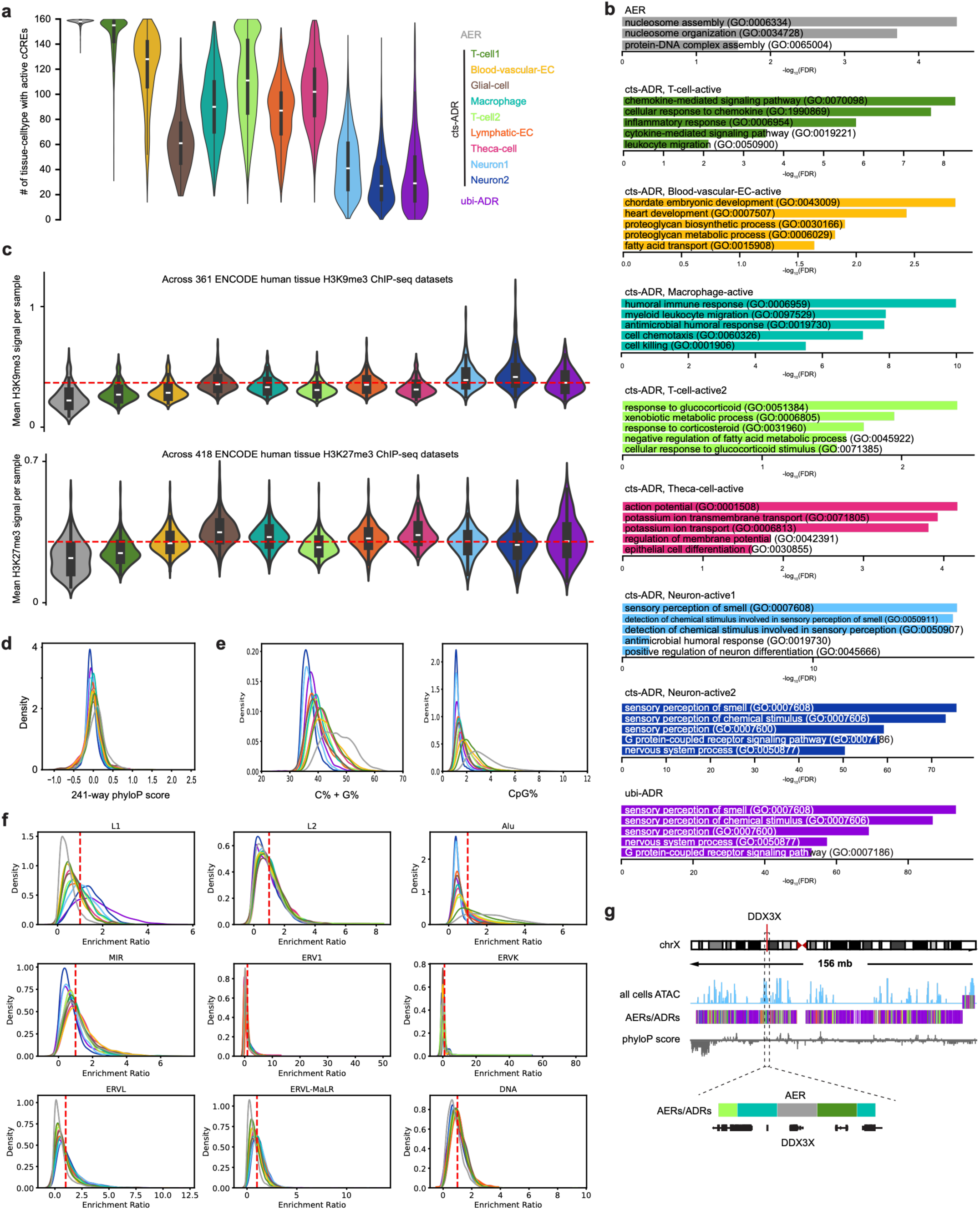
Genomic and epigenomic features of broad chromatin-domain classes. **a.** Distribution of the number of cell clusters in which each 100-kb bin is ATAC-depleted, stratified by broad chromatin-domain class. Colors indicate cts-ADR subclasses and ubi-ADR subclasses. **b.** Gene Ontology enrichment analysis for representative domain classes. Bars show selected enriched terms for AERs, T-cell-specific cts-ADRs, blood-vascular-endothelial-cell-specific cts-ADRs, macrophage-specific cts-ADRs, T-cell-specific cts-ADRs, theca-cell-specific cts-ADRs, neuron-specific ubi-ADRs and general ubi-ADRs. **c.** Distribution of H3K9me3 and H3K27me3 ChIP-seq signal across broad chromatin-domain classes, based on ENCODE human tissue datasets. Violin plots show the mean signal per genomic bin; the dashed red line indicates the genome-wide reference level. **d.** Density distributions of 241-way placental mammal phyloP scores across genomic bins, stratified by domain class. **e.** Density distributions of total GC content and CpG dinucleotide content across genomic bins, stratified by domain class. **f**. Density distributions of observed-to-expected enrichment ratios for major transposable-element families across domain classes. The dashed red line indicates an enrichment ratio of 1. **g**. Genome-browser views of chrX illustrating broad chromatin-domain classes. Tracks show ATAC-seq signal across all cells, together with AERs, ADRs and phyloP conservation scores.

**Extended Data Fig. 8.**
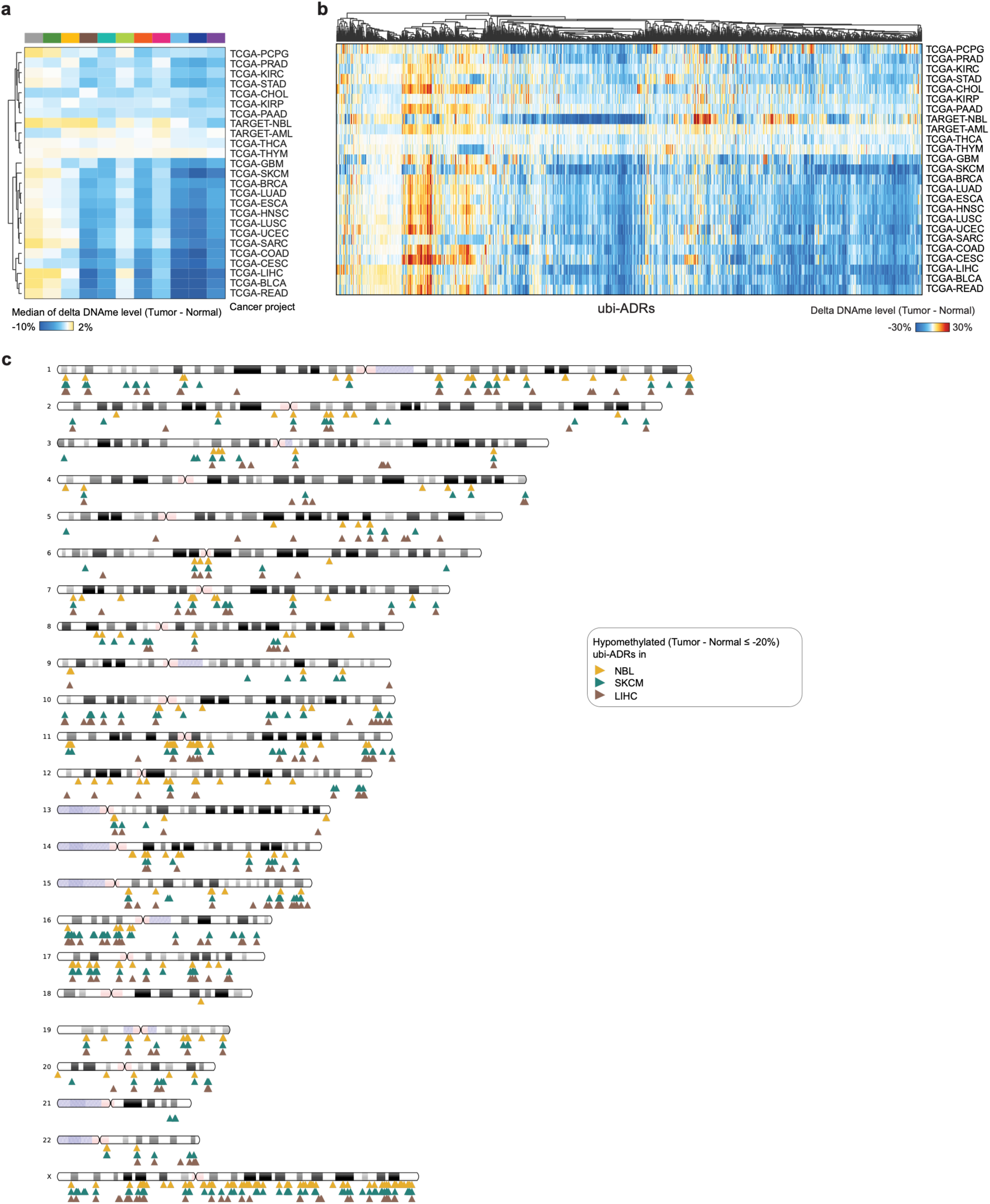
Tumour-associated DNA methylation changes across broad chromatin domains. **a**. Heatmap showing the median change in DNA methylation between primary tumours and matched normal tissues across broad chromatin-domain classes. Rows represent TCGA and TARGET cancer projects, and columns represent domain classes. Colors indicate the median difference in DNA methylation (tumour minus normal). TCGA-LIHC, liver hepatocellular carcinoma; TCGA-SKCM, skin cutaneous melanoma; TCGA-BLCA, bladder urothelial carcinoma; TCGA-READ, rectum adenocarcinoma; TCGA-SARC, sarcoma; TCGA-COAD, colon adenocarcinoma; TCGA-CESC, cervical squamous cell carcinoma and endocervical adenocarcinoma; TCGA-BRCA, breast invasive carcinoma; TCGA-HNSC, head and neck squamous cell carcinoma; TCGA-UCEC, uterine corpus endometrial carcinoma; TCGA-LUSC, lung squamous cell carcinoma; TCGA-ESCA, oesophageal carcinoma; TCGA-LUAD, lung adenocarcinoma; TCGA-GBM, glioblastoma multiforme; TCGA-PRAD, prostate adenocarcinoma; TCGA-STAD, stomach adenocarcinoma; TARGET-NBL, neuroblastoma; TCGA-CHOL, cholangiocarcinoma; TCGA-KIRP, kidney renal papillary cell carcinoma; TCGA-PCPG, pheochromocytoma and paraganglioma; TCGA-KIRC, kidney renal clear cell carcinoma; TCGA-PAAD, pancreatic adenocarcinoma; TCGA-THCA, thyroid carcinoma; TCGA-THYM, thymoma; and TARGET-AML, acute myeloid leukaemia. **b**. Heatmaps showing changes in DNA methylation between primary tumours and matched normal tissues across genomic bins within ubi-ADRs. Rows represent TCGA and TARGET tumour cohorts, and columns represent genomic bins ordered by hierarchical clustering. Colors indicate the change in DNA methylation (tumour minus normal), with blue denoting relative hypomethylation and orange denoting relative hypermethylation in tumours. **c**. Chromosomal distribution of ubi-ADRs hypomethylated in neuroblastoma (TARGET-NBL), skin cutaneous melanoma (TCGA-SKCM) and liver hepatocellular carcinoma (TCGA-LIHC). Selected genes overlapping or located near hypomethylated ubi-ADRs are indicated.

**Extended Data Fig. 9.**
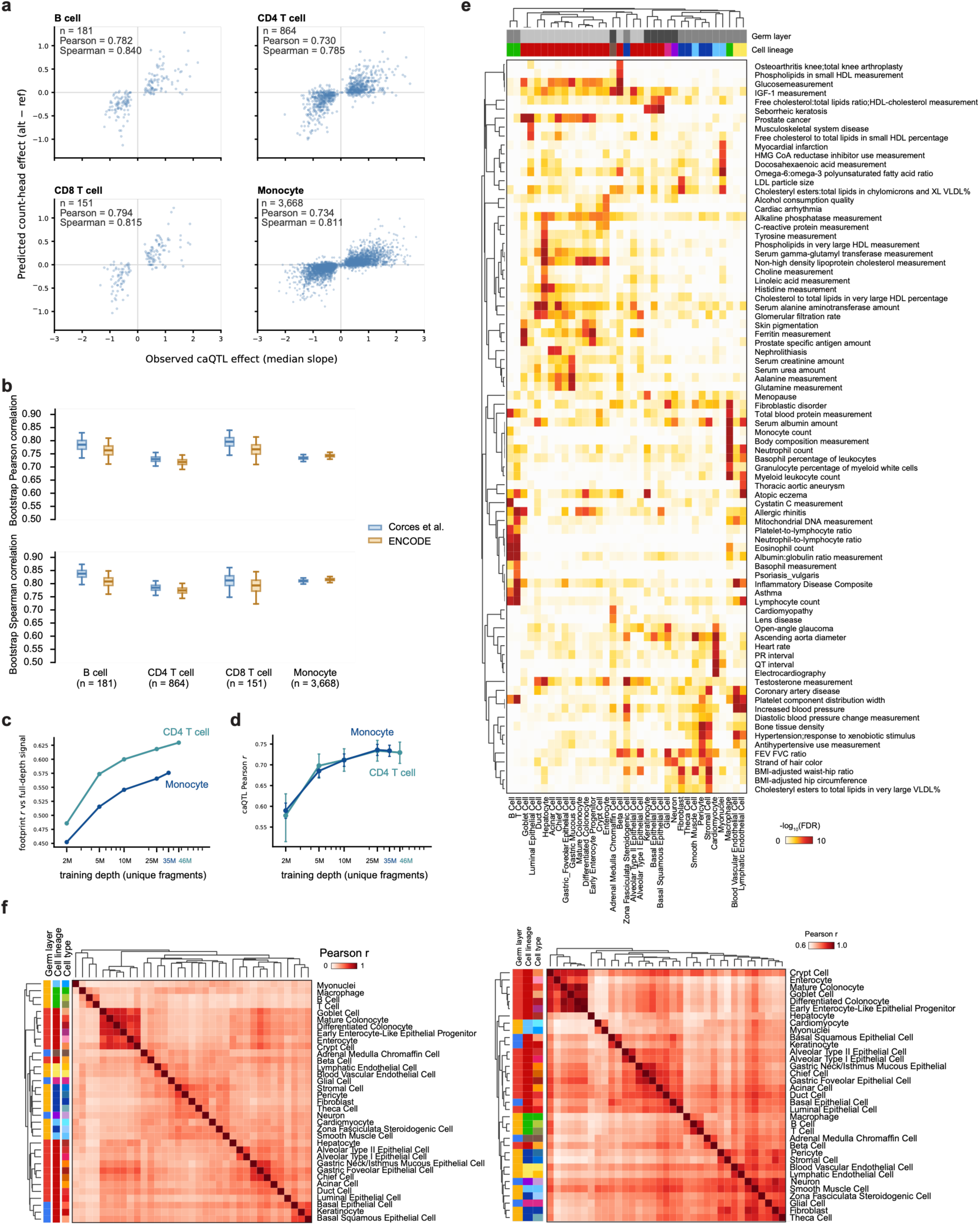
Cell-type-resolved regulatory effects of complex-trait-associated variants. **a.** Predicted regulatory effects plotted against observed caQTL effects for B cells, CD4 T cells, CD8 T cells and monocytes using data from Corces *et al*. **b.** Bootstrap Pearson correlations between predicted effects and caQTL effects for models trained on Corces *et al*. and ENCODE data. Box plots show the distribution of bootstrap correlations. **c.** Pearson correlations between predicted regulatory effects from seq2PRINT and full-depth ATAC-seq shown as a function of sequencing depth for CD4 T cells and monocytes. **d**. Pearson correlations between predicted regulatory effects from seq2PRINT and caQTL shown as a function of sequencing depth for CD4 T cells and monocytes. Error bars indicate 95% bootstrap confidence intervals. **e**. Heatmap showing associations between GWAS studies spanning diverse trait categories and the 35 primary cell types. Traits are grouped by broad phenotype category. Colors indicate −log_10_(FDR), and annotation bars indicate germ layer, cell lineage and cell-type category. **f**. Pairwise Pearson correlations between seq2PRINT-predicted regulatory-effect profiles (left) and ATAC signal (right) across the 35 primary cell types. Rows and columns are ordered by hierarchical clustering. Annotation bars indicate germ layer, cell lineage and cell type.

**Extended Data Fig. 10.**
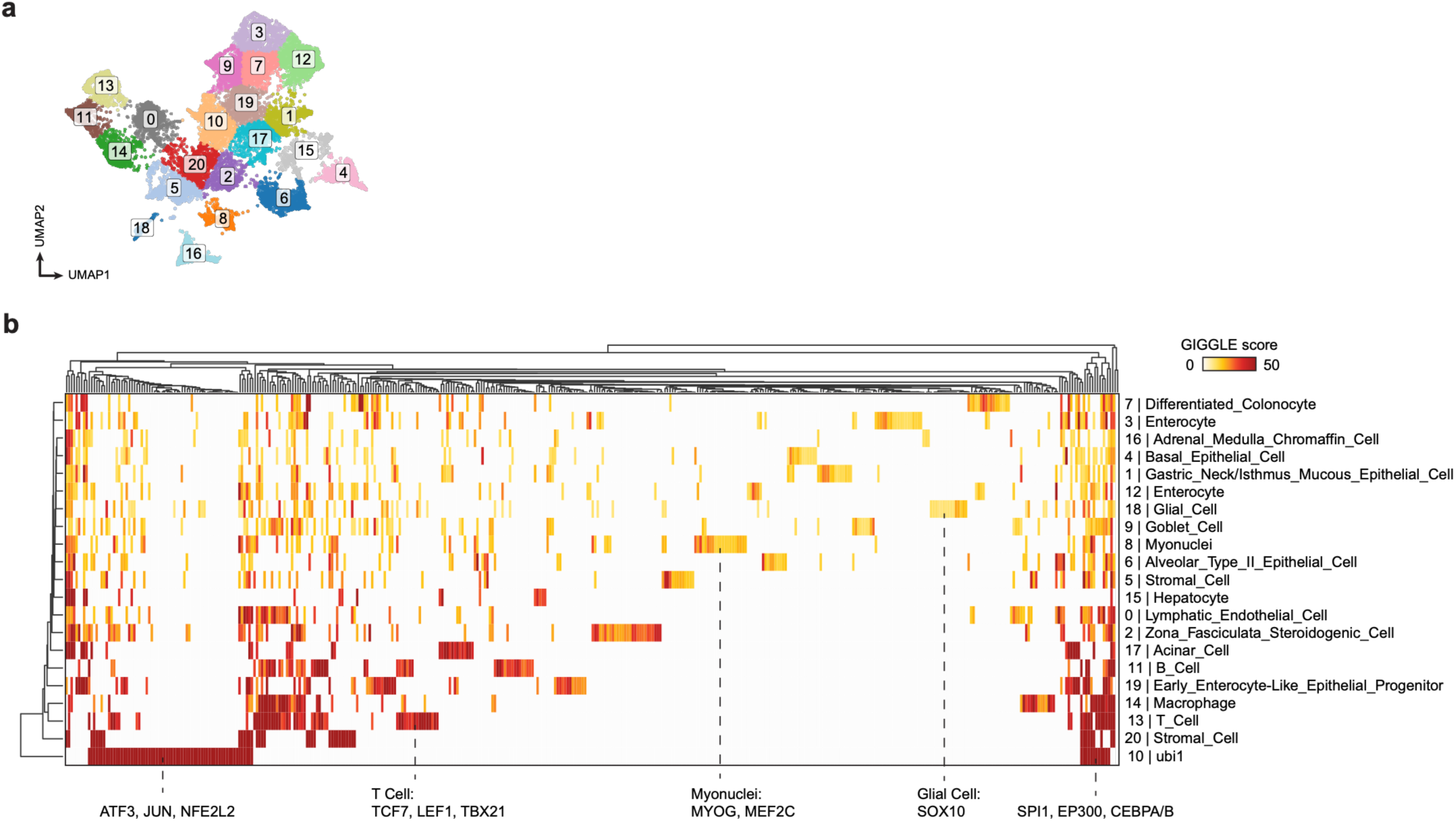
Diverse TF binding across different cluster of pcaHEVs. **a.** UMAP embedding of pcaHEVs colored by the cell type in which each variant shows the highest predicted regulatory effect. **b.** Heatmap of GIGGLE scores comparing pcaHEV-associated cell-type profiles with transcription-factor cistrome datasets. Rows represent transcription factors and columns represent cell types ordered by hierarchical clustering. Colors indicate the GIGGLE score.

